# Visualizing and manipulating hyperstabilization of T cell microvilli contacts by engineered chimeric antigen receptors

**DOI:** 10.1101/2021.08.18.456686

**Authors:** Casey Beppler, John Eichorst, Kyle Marchuk, En Cai, Carlos A. Castellanos, Venkataraman Sriram, Kole T. Roybal, Matthew F. Krummel

**Author notes:** These authors contributed equally (Kole T. Roybal, Matthew F. Krummel).

## Abstract

T cells typically recognize their ligands using a defined cell biology – the scanning of their membrane microvilli to palpate their environment – while that same membrane scaffolds T cell receptors (TCRs) that can signal upon ligand binding. Chimeric antigen receptors (CARs) present both a therapeutic promise as well as a tractable means to study the interplay between receptor affinity, microvillar dynamics and T cell function. CARs are often built using single-chain variable fragments (scFvs) with far greater affinity than that of natural TCRs. We used high resolution lattice lightsheet (LLS) and total internal reflection fluorescence (TIRF) imaging to visualize microvillar scanning in the context of variations in CAR design. This demonstrated that conventional CARs hyper-stabilized microvillar contacts relative to TCRs. Reducing the affinity and/or avidity of binding brought synapse microvillar dynamics into natural ranges, normalized synapse resolution and improved downstream effector function. This work highlights the importance of understanding the underlying cell biology when designing receptors for optimal antigen engagement.

## Introduction

While epithelial cell microvilli are historically well characterized for their cell biology (Crawley et al., 2014; Sauvanet et al., 2015), advancements in microscopy have shown dynamic microvilli on T cells (Majstoravich et al., 2004) to be critical for the cell’s antigen search process (Cai et al., 2017; Orbach and Su, 2020; Pettmann et al., 2018). During the antigen scanning phase of early immunological synapse formation, microvilli enable close contact with antigen-presenting surfaces and become stabilized specifically upon recognition of cognate antigen (Cai et al., 2017). Furthermore, microvilli have been shown to contribute to T cell signaling by promoting optimal kinetic segregation of phosphatases away from signaling domains (Razvag et al., 2018), and inducing the formation and confinement of microvilli has even been shown to be sufficient for T cell activation in the absence of TCR agonists (Aramesh et al., 2021).

T cell recognition occurs at immune synapses (IS), which are well studied in two dimensions on supported lipid bilayers (SLBs). TCR signaling at the prototypical immune synapse involves the local accumulation of signaling components colocalizing with microclusters of TCRs. After a few minutes, these can internalize (Friedl et al., 2005; Friedman et al., 2010) and/or migrate toward the center of the synapse into canonical central supramolecular activation clusters (cSMAC) (Campi et al., 2005; Grakoui et al., 1999; Monks et al., 1998). The cSMAC – where active kinases are rare – is a site where TCR signaling is thought to end (Campi et al., 2005; Varma et al., 2006). Microclusters are thought to be conducive to signal initiation and amplification – allowing two-dimensional segregation of phosphatases away from signaling domains. More recent work has revealed that TCR microclusters form on the tips of dynamically probing microvilli, facilitating their stabilization, and it is upon these microvillar protrusions that microclusters move laterally within the early synaptic interface (Cai et al., 2017).

Chimeric antigen receptor-bearing T cells (CAR T cells) present both a therapeutic promise, given their success in the treatment of B cell malignancies (Kalos et al., 2011; Brentjens et al., 2013; Maude et al., 2014; Turtle et al., 2016), and a tractable system for understanding how affinity and avidity affect topographical synapse antigen scanning, given that they are easily engineered molecules. Typical CAR designs used for therapy incorporate a tumor antigen-binding scFv, along with a hinge/transmembrane domain (commonly from either CD8 or CD28) and intracellular CD3ζ and co-stimulatory domains (commonly from either 4-1BB or CD28). Signaling events initiated by scFv binding and subsequent phosphorylation of intracellular components at the immune synapse lead to the acquisition of CAR T cell effector functions. Prior to this work, a recent study demonstrated aberrant synapse architecture formed by T cells downstream of CAR binding (Davenport et al., 2018), although factors leading to that synaptic phenotype were not obvious. Whether and how CARs engage the underlying native cell biology remain largely unknown.

The antigen binding domains of engineered CARs and the T cell’s natural antigen receptor differ significantly. CAR scFvs bind tumor antigen with nano- and even picomolar affinities whereas TCRs bind agonist peptide-major histocompatibility complex (pMHC) with micromolar affinity (Stone et al., 2009), although functional binding strength is improved by contributions of co-receptor (Garcia et al., 1996). The effect of such variation in binding on the dynamics of topographical antigen scanning is unknown. This is an especially interesting question given that CARs generate inefficient intracellular signaling relative to TCRs (Harris et al., 2018; Gudipati et al., 2020; Salter et al., 2021). It therefore seems plausible that improving the already ultra-high affinity binding might be beneficial. On the other hand, it is well accepted that T cells achieve high sensitivity through serial engagement (Viola et al., 1997; Hudrisier et al., 1998) and thus the T cell’s biology may be tuned to respond to a specific range of receptor binding dynamics (Corse et al., 2010; McMahan et al., 2006; Kalergis et al., 2001; Schmid et al., 2010; Hebeisen et al., 2013).

Recent work shows that even within natural TCR affinity ranges, there is a “sweet spot” for binding strength (Shakiba et al., 2021). We thus sought to understand how CARs interact with the natural cell biology, relative to TCRs, and how varying the binding dynamics of an antigen receptor affects topographical antigen scanning.

To address these questions, we combined high-resolution lattice lightsheet (LLS) and synaptic contact mapping (SCM) total internal reflection fluorescence (TIRF) imaging – first, to establish the distribution of CAR relative to TCR on the T cell surface both prior to and during antigen detection, and second, to establish the effects of receptor affinity and avidity on the dynamics of topographical scanning. In order to study how topographical scanning dynamics are affected by receptor affinity and avidity in isolation, point mutations were incorporated into the extracellular domains of the engineered receptor while all other aspects of CAR design were held constant – hinge and transmembrane domain (CD8), co-stimulatory domain (4-1BB), and CD3ζ signaling domain. In doing so, we found that CARs distributed similarly to TCRs. However, CARs based on conventional nanomolar high affinity dimers yielded hyperstabilization of the underlying microvillus relative to that of the natural TCR, which could be improved by reducing receptor affinity or avidity. Hyperstabilization was correlated with aberrant synapse resolution, decreased effector function, and increased propensity for exhaustion.

## Results

### TCR and CAR distribution relative to microvilli on isolated T cells

TCRs have been reported to form patches (Lillemeier et al., 2006; Hu et al., 2016; Cai et al., 2022) and localize to microvilli to varying degrees (Jung et al., 2016; Cai et al., 2017, 2022). We expressed anti-HER2 CARs in primary human CD8+ T cells by lentiviral transduction and first sought to compare how they distribute on the T cell surface, compared to conventional TCR. To do this, we first fixed CAR T cells, stained them with antibodies that bound to TCR and CAR, and imaged them by LLS. Visually, this revealed that CARs were distributed across the surface of the cell with patches of increased fluorescence intensity, similar to that of TCR (Cai et al., 2022) although not necessarily overlapping in position (**Fig. 1a-c, Supplemental Fig. 1a-b** and **Supplemental Video 1**). Although examined here on fixed cells, similar patches have also been described for TCRs by live imaging (Cai et al., 2022). Zooming in to a region of the cell membrane, microvilli were thus noted as variably high for CAR alone, TCR alone, or both (**Fig. 1b** and **Supplemental Fig. 1c**). About half of microvilli tips observed were high for TCR (either with or without CAR), and most but not all (83%) microvilli tips were CAR high (**Fig. 1c** and **Supplemental Fig. 1d**), likely resulting from high CAR expression induced by lentiviral expression. Random distribution of CAR and TCR based off these proportions would predict ∼40% co-occupied tips. By our count, 32% of tips are co-occupied, consistent with the co-occupancy being a random event.

**Figure 1:**
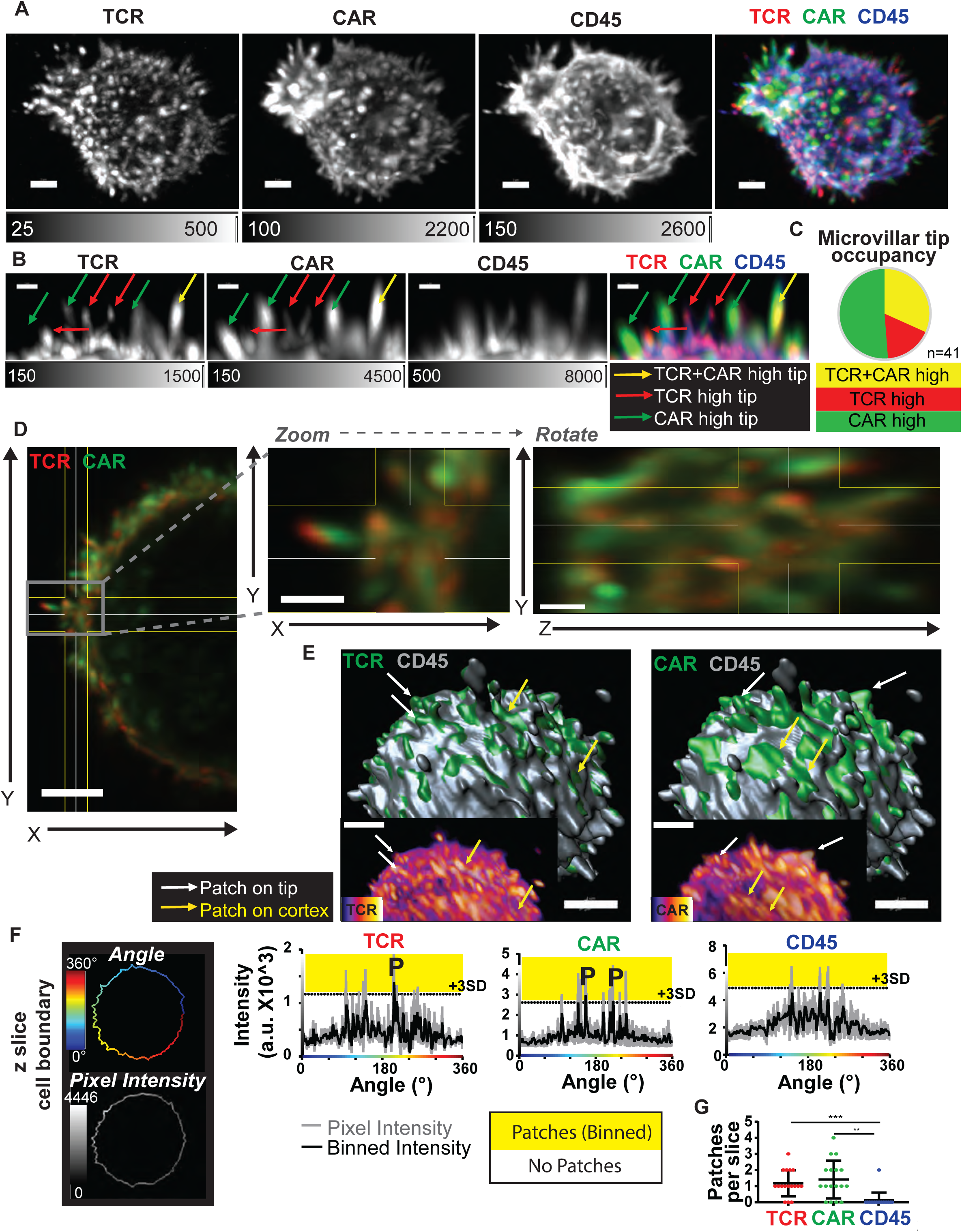
CARs distribute similarly to TCRs on both microvilli and cell cortex, but do not obligately co-localize. A. Fixed anti-HER2 CAR T cell showing maximum intensity projection of TCR, CAR, CD45 and overlay imaged by lattice light-sheet. Scale bar = 2 μm. Human CD8+ T cell expressing anti- HER2 CAR (mut4D5) was labelled with antibodies to MYC (CAR)-Alexa488, TCR (OKT3)-APC, and CD45-Alexa594. B. Region of cell membrane, as described in A, where microvilli tips are marked as being high for TCR (red arrow), CAR (green arrow), or both (yellow arrow). Scale bar = 1 μm. C. 41 tips across 5 regions and 3 cells were scored as in B. D. Left panel shows zoomed out membrane region in extended view for multiple z slices of a cell labelled as described in A (scale bar = 3 μm). Middle panel shows zoom in on outlined region (scale bar = 1 μm). Right panel shows the same zoom region with the perspective rotated to show the YZ plane (scale bar = 1 μm). E. Large panels show CD45 surface (gray) overlaid with TCR (left) and CAR (right) surfaces marking areas of high intensity (green). Arrows mark examples of patches on microvilli tips (white) or cell cortex (yellow). Inset panels show a region of the same cell with the raw TCR and CAR intensity in Fire colorscale. Scale bar = 4 μm. F. Left: Schematic showing the cell boundary as mapped to radial intensity profile by angle (x- axis) and pixel intensity (y-axis). Right: Pixel intensity (gray line) and binned intensity (black line), defined by taking a moving average of 10 pixels, showing one z-slice for TCR, CAR, and CD45. Dashed line with yellow above indicates 3 standard deviations (+3SD) above mean pixel intensity. P indicates binned intensity peaks above +3SD threshold (patches). G. Number of patches/z-slice for each receptor was defined by the number of excursions of binned intensity above the +3SD threshold. Tukey’s multiple comparisons test was performed on data compiled from multiple slices of 3 cells (n = 17 slices per group). Error bar represents standard deviation (s.d.).

Of note, not all CAR or TCR patches occurred on microvilli – TCR high, CAR high, and TCR/CAR high patch-like regions could be seen in both tips and areas of cell cortex, here referring to the regions on the body of the cell between microvilli (**Fig. 1D**). This was further demonstrated by overlaying the surface of the T cell based off CD45 expression with surfaces of high intensity TCR and CAR “patches” (**Fig. 1e**). To quantify this patchiness, the fluorescence intensity at the membrane was mapped to radial intensity profiles (**Fig. 1f** and **Supplemental Fig. 1e**). Taking a moving average with a bin size of 10 pixels (∼3° degrees), patches of increased local fluorescence were identified for TCRs, as expected(Hu et al., 2016), but less so for the abundant CD45 molecule. Using this method, CARs were found to form patches at frequencies similar to TCRs (**Fig. 1g**) despite not obligately co-localizing.

In order to assess TCR and CAR patches relative to surface curvature in three dimensions, membrane surface curvature was mapped based on CD45 (**Fig. 2a** and **Supplemental Fig. 1f**) and regions of high convexity were thresholded to show microvilli (**Fig. 2b**). CAR and TCR patches were assigned by identifying individual domains of locally high intensity along the cell boundary using Pearson correlation calculations for variable sized kernels followed by segmentation and watershed (Cai et al., 2022) (**Fig. 2c-d** and **Supplemental Fig. 1f**). Surface areas of individual patches were then mapped by color (**Fig. 2e-f**). A histogram overlay and dot plots of hundreds of patches compiled from three cells show similar distribution of patch surface area for TCR and CAR, with median patch sizes of 0.34 μm^2^ (TCR) and 0.38 μm^2^ (CAR) (**Fig. 2g**). Surface curvature values on cells were mean-centered and both TCR and CAR patch localization were similarly mean-centered, indicating that there is not a preference for patches to reside in regions of high or low surface curvature values (**Fig. 2h**). Two-dimensional projections of the three-dimensional surface curvature and antigen receptor intensity were made and overlaid, revealing regions of membrane tips and valleys both with and without antigen receptor (**Fig. 2i**). Relative co-localization of TCR and CAR patches was assessed by counting the number of TCR and CAR patches that met varying thresholds of overlap with CAR and TCR patches, respectively. Total number of identified patches (0% colocalization threshold) was similar for CAR and TCR in a given cell, and the majority of patches for each receptor had <50% overlap with clusters of the other (**Supplemental Fig. 1g-h**).

**Figure 2:**
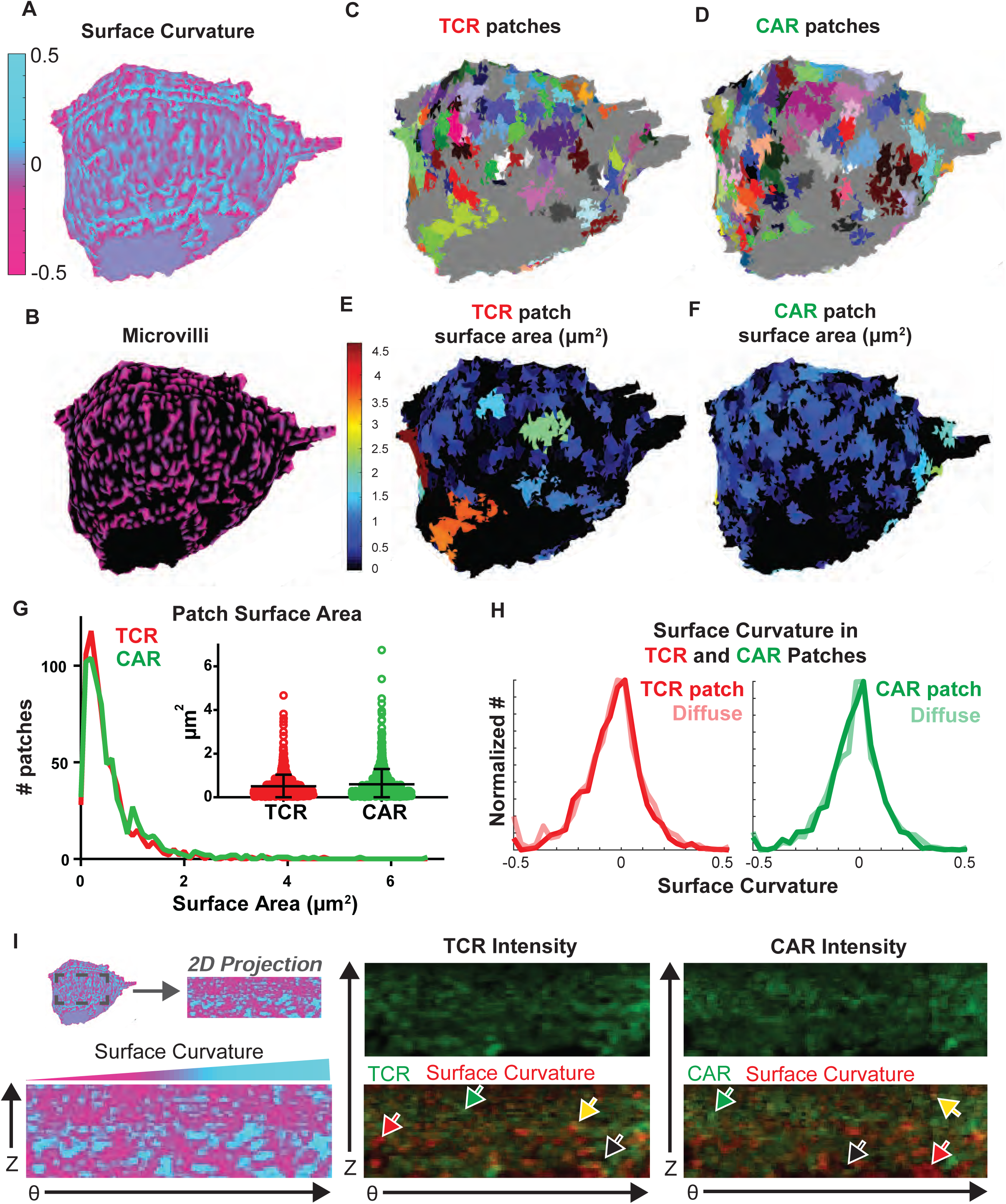
Quantification of TCR and CAR patches relative to surface curvature. A. Curvature of the cell surface was mapped based off the cell boundary defined by CD45, with valleys marked by cyan and tips marked by magenta. Fixed CAR T cell was labelled with antibodies to MYC (CAR)-Alexa488, TCR (OKT3)-APC, and CD45-Alexa594. B. Microvilli are highlighted by masking the surface curvature in A based on convexity. C-F. TCR (C, E) and CAR (D, F) clusters defined by regions of high local (3D) intensity are shown as different colors (C-D) and colored according to surface area of cluster (E-F). G. Histogram and dot plot (inset) of cluster surface area is shown for TCR and CAR. TCR and CAR median values are 0.34 and 0.38 μm^2^, respectively. Data is compiled from three cells. Dot plot shows mean and s.d. H. Histograms of surface curvature for TCR (left) and CAR (right) found in patches or outside of patches. I. 2D projections from the surface show curvature increasing from convex (magenta on left, black in overlay) to concave (cyan on left, red in overlay); TCR intensity (middle); and CAR intensity. Yellow and red arrows mark membrane valleys with high and low receptor intensity, respectively. Green and black arrows mark membrane peaks with high and low receptor intensity, respectively.

### CAR enrichment at synaptic microvillar close contacts

Next, in order to determine the location of CARs following interaction with a target cell, live anti- HER2 CAR T cells were added to coverslips loaded with HER2+ SKBR3 cells and imaged by LLS at intervals of 4.7 (**Fig. 3a-c, Supplemental Fig. 2a** and **Supplemental Video 2**) or 6.75 (**Supplemental Fig. 2b**) secs. This live cell imaging revealed accumulation of CAR patches at projections into the synapse.

**Figure 3:**
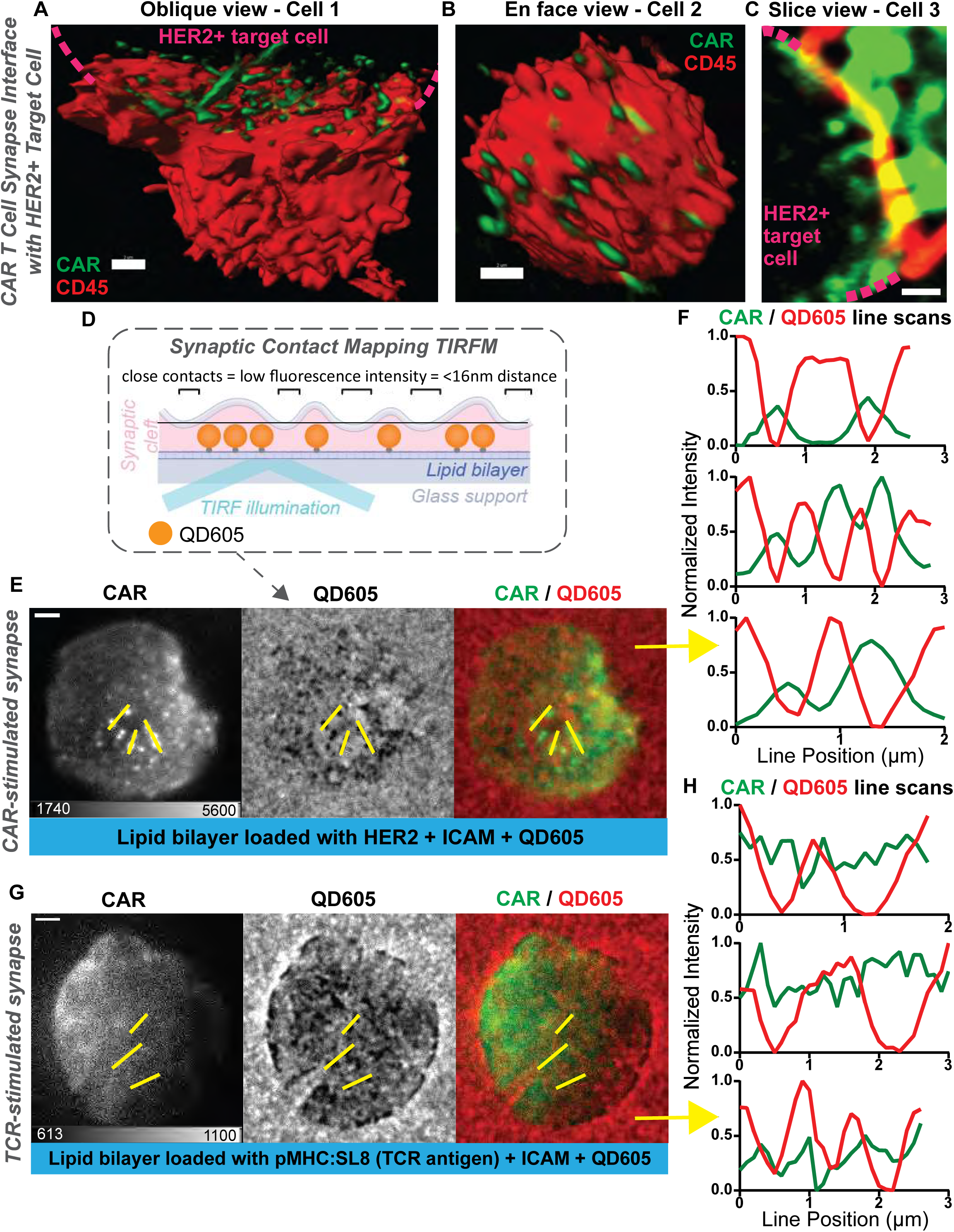
CAR is enriched at sites of microvilli close contacts in synapses with cognate antigen, but not following engagement of endogenous TCR. A-C. Three examples of anti-HER2 CAR T cells interacting with HER2+ SKBR3 are shown from an oblique view of the synapse interface (A, Imaris blended view, scale bar = 2 μm), en face synapse view (B, Imaris blended view, scale bar = 2 μm), and a single z-slice (C, scale bar = 1 μm). The SKBR3 target cell location is marked by dashed magenta lines (A, C). Human CD8+ T cell expressing anti-HER2 CAR (4D5) was labelled with antibodies to MYC (CAR)-Alexa488 and CD45-Alexa647. D. Schematic of synaptic contact mapping method. Lipid bilayer on glass support is loaded with Qdot 605, which is a red-fluorescent probe that occupies ∼16nm in height. Areas where the cell makes a close contact (<16nm distance) appear as holes in the QD605 signal. E. CAR T cell synapse imaged by TIRF showing CAR-mEmerald, microvillar projections (as seen by holes in QD605 signal), and overlay. Human CD8+ T cell expressing anti-HER2 CAR (mut4D5) interacting with lipid bilayer loaded with 625ng HER2 + ICAM + QD605. Scale bar = 2 μm. F. Line scans from F show anti-correlation of CAR-mEmerald and QD605, indicating enrichment of CARs within microvillar contacts. G. OT-I mouse T cells were retrovirally transduced with anti-HER2 CAR (mutCD45). Synapse shown was formed on lipid bilayer loaded with pMHC:SL8 + ICAM + QD605. TIRF imaging of CAR-mEmerald, QD605, and overlay are shown. Scale bar = 2 μm. H. Line scans from H showing the lack of enrichment in CAR signal within QD605 holes, indicating that CAR microclusters do not accumulate in microvillar close contacts in absence of the CAR’s cognate antigen.

To further study the localization of CARs to these microvillar close contacts, mEmerald-tagged CAR T cells were analyzed on HER2-loaded lipid bilayers using SCM TIRF microscopy. This method allows for facile visualization of both the density of receptors near the surface by TIRF in a green fluorescent channel, and the location of microvillar close contacts, as seen by holes in fluorescent quantum dot (QD605) signal (Cai et al., 2017) (**Fig. 3d**). As previously demonstrated for TCR (Cai et al., 2017), microclusters of CAR-mEmerald fluorescence were found to form and localize to areas of microvillar close contacts, as seen by line scans revealing segregation between the two signals (**Fig. 3e-f** and **Supplemental Fig. 2d,f**). As with TCR, not all microvillar close contacts at the synapse were occupied by CAR microclusters (**Supplemental Fig. 2c**). To determine whether this localization was dependent simply on the formation of a stabilized close-contact or depended specifically on binding of CAR to its ligand, anti-HER2 CARs were expressed in OT-I mouse T cells and allowed to interact with bilayers loaded with only the TCR ligand, pMHC:SL8 (SIINFEKL). In these HER2-negative synapses microvillar contacts were still formed but CARs did not accumulate as microclusters nor localize within those contacts (**Fig. 3g-h** and **Supplemental Fig. 2e,g**). Thus, we found that the recruitment of CAR microclusters to the microvillar close contacts relied upon the binding of CAR antigen on the interacting surface, and that TCR engagement was not sufficient to coordinate the CAR’s reorganization at the interface.

### Effects of affinity and avidity of receptor binding on synaptic microvillar dynamics

We next sought to understand how microvillar close contacts, hallmarks of TCR-induced activation (Cai et al., 2017; Farrell et al., 2020), were modulated by the affinity of the antigen receptor. To do this, we compared the wild-type high-affinity scFv (4D5) with a mutated scFv of ∼70-fold reduced affinity (mut4D5, **Fig. 4a** and **Supplementary Table 1**). We then analyzed microvillar close contacts by SCM at high temporal resolution in the presence of a range of ligand densities to measure microvillar persistence times (**Fig. 4b-d**). Ligand densities were standardized against two breast cancer cell lines that differed significantly in their HER2 levels, and addition of 62.5 ng per 0.7 cm^2^ well yields expression levels comparable to a cell line representing healthy breast tissue, MCF7 (Nobili et al., 2021; Liu et al., 2015; Comsa et al., 2015) (**Supplemental Fig. 2h-i**). In the examples in **Fig. 4c** and **Supplemental Video 3,** individual microvillar close contacts mediated by low-HER2 bilayers remained stable in the case of a high-affinity CAR contact, while the low-affinity CAR close contact dispersed within seconds. The lifetime of individual microvillar close contacts was measured using automated scripts that identify regions of QD605-exclusion. This revealed that the persistence time of CAR-occupied close contacts increases with ligand density for low affinity CARs, but that high affinity CAR resulted in very stable contacts at all tested ligand densities (**Fig. 4d**). All conditions tested induced stabilization of microvilli above background non-receptor-occupied contacts, which had uniformly short persistence times. Notably, at high HER2 density, the persistence of low-affinity CAR close contacts reached ∼17 secs on average, a value that was achieved by high-affinity CAR even at 100x lower antigen densities – lower even than the low-HER2 MCF7 standard. Importantly, both low (62.5 ng/well) and high (625 ng/well) antigen densities led to calcium flux for both receptors, as detected by Fura-2 ratiometric imaging (**Supplemental Fig. 2j-k** and **Supplemental Video 4**). At high density of antigen, calcium imaging demonstrated a decrease in the magnitude of the maximal ratio achieved, specifically for the high-affinity CAR (**Supplemental Fig. 2k**), suggesting altered signaling dynamics in the high-affinity setting.

**Figure 4:**
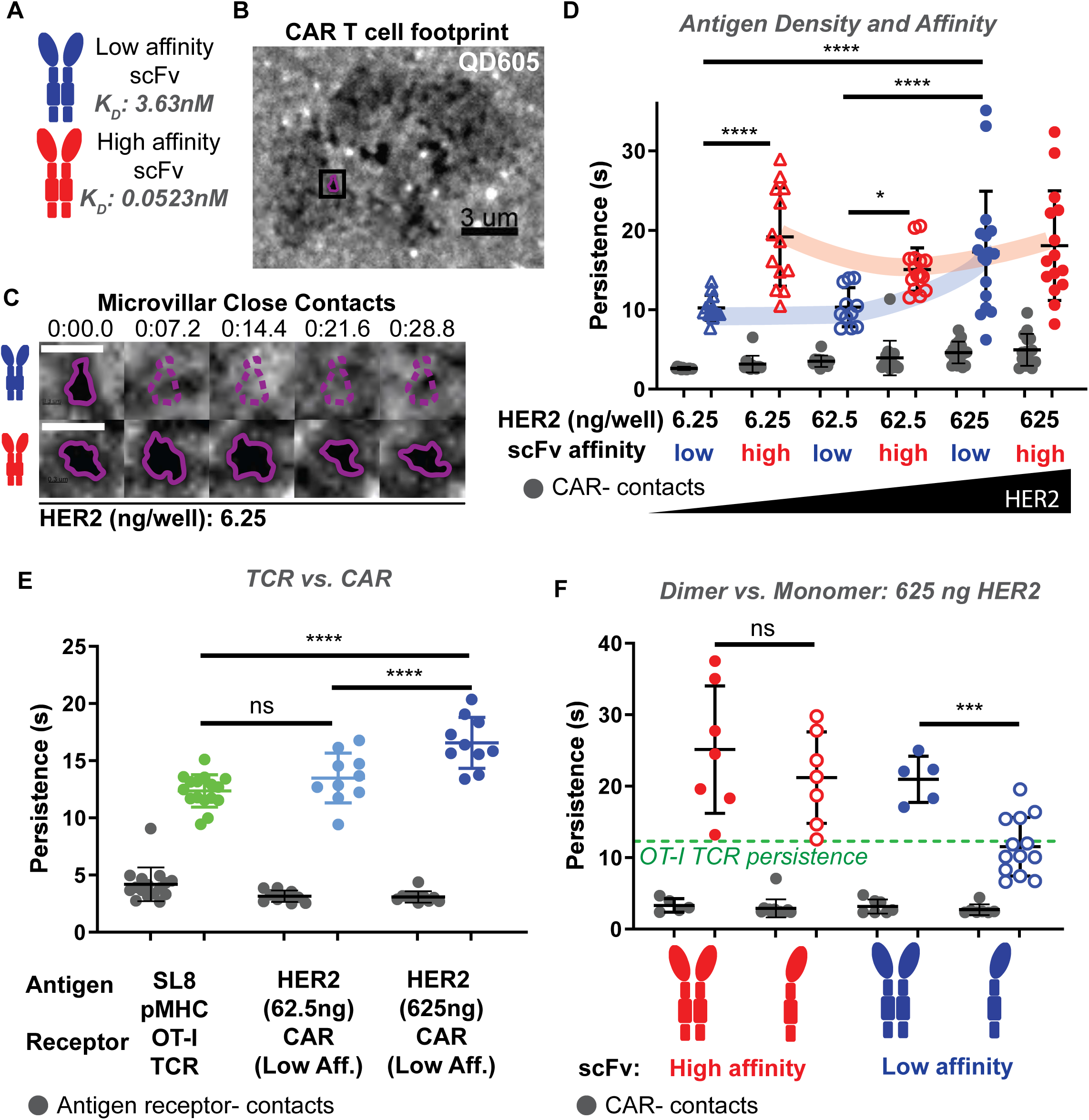
Conventional CAR interactions of high affinity or high antigen density result in hyper-stabilization of underlying microvillar protrusion, which can be reduced by using monomeric CAR. A. Experiments were performed comparing CARs with a high-affinity scFv against HER2-based off trastuzumab (4D5, KD = 0.0523nM, red) and a lower affinity scFv made by substitution of three amino acids (see Supplemental Table 1, mut4D5, KD = 3.63nM, blue). B. Bilayers were loaded with fluorescent quantum dots (QD605) with a height of 16nm. Locations where the cell makes close contact with the bilayer (<16nm) are visualized as holes in QD605 signal due to their size-based exclusion. QD605 signal is shown for a low-affinity CAR T cell interacting with a lipid bilayer loaded with 6.25 ng HER2. Outlined contact is shown in C. Scale bar = 3 μm. C. QD605 signal across 5 time points are shown for the same field of view, each on low HER2 bilayers (6.25 ng/well). Top: Microvillus from low-affinity CAR T cell moves out of view. Bottom: Microvillus from high-affinity CAR T cell remains across time points. White scale bar = 1 μm. D. CAR-occupied close contact persistence times (blue, red) and CAR-negative close contact persistence times (gray) are shown for varying antigen densities and affinities. All CAR:HER2 interactions tested result in CAR-occupied microvillar contact stabilization above background CAR-negative contacts (gray). Persistence time is further increased in interactions of high- affinity CAR (red), even at lowest HER2 densities on the bilayer. For low-affinity CAR (blue), only high levels of HER2 yield similar persistence times to high-affinity CAR. Data is shown for at least 11 cells per condition across four experiments (n = 84, 13, 13, 11, 15, 17, 15 cells per group from left to right, respectively). E. Low-affinity CAR was retrovirally expressed in primary mouse OT-I T cells. Receptor- occupied microvillar persistence times are shown for OT-I:SL8 (green), Low-affinity CAR:Low HER2 (light blue), and Low-affinity CAR:High HER2 (dark blue) interactions. All cognate interactions are stabilized above background receptor-negative contacts (gray). CAR:High HER2 persistence is hyper-stable relative to TCR:pMHC stabilization. Data is shown for at least 10 cells per condition across three experiments (n = 36, 16, 10, 10 cells per group from left to right, respectively). F. Dimers (filled dots) and monomers (open dots) are compared on high HER2 bilayers (625ng/well). Only monomeric low-affinity CAR regains natural persistence time of TCR:pMHC contacts (green dashed line). All receptor-occupied contacts are stabilized above non-cognate antigen interactions (gray). Data is shown for at least 5 cells per condition across three experiments (n = 32, 7, 7, 5, 13 cells per group from left to right, respectively). All error bars represent s.d. and analyses shown are Šídák’s multiple comparisons tests.

We next sought to directly compare CAR- and TCR-mediated persistence times. We thus expressed the low-affinity CAR in OT-I mouse T cells and compared interactions with SL8:pMHC- and HER2-loaded bilayers. On bilayers with SL8:pMHC, normalized to levels displayed on APCs (Beemiller et al., 2012a), we measured TCR-mediated persistence to be an average of 12.3 seconds. Low-affinity CAR with low-HER2 bilayers yielded similar persistence times whereas persistence times on high HER2 were again found to be significantly greater (**Fig. 4e**). We hypothesized that we could further engineer the low-affinity CAR to yield TCR-like persistence times in high antigen-density conditions by minimizing CAR avidity contributions. Thus, we created a monomeric CAR by incorporating two cysteine to serine point mutations in the CD8α hinge to prevent formation of the disulfide bridge (Hennecke and Cosson, 1993) (**Supplemental Fig. 2l**). Comparing the persistence time of monomers to dimers on high HER2-loaded bilayers (625ng/well) revealed that only the low-affinity monomers produced microvillar persistence similar to that of the TCR (**Fig. 4f**). Average CAR fluorescence intensity of close contacts was not significantly different between monomers and dimers and did not appear to correlate with persistence times under the conditions tested (**Supplemental Fig. 2m-n**).

### Hyperstable microvillar dynamics are associated with altered synapse resolution, decreased effector function, and increased propensity for exhaustion

TCR-mediated synapses formed on antigen-loaded lipid bilayers result in TCR microcluster formation and radial motion of microclusters toward the center (Campi et al., 2005; Yokosuka et al., 2005; Varma et al., 2006). Although alternative synapse architectures characterized by the formation of CAR microclusters without centripetal movement or cSMAC formation have been revealed for CAR-mediated synapses (Davenport et al., 2018), we sought to relate binding dynamics resulting in hyperstabilization of microvilli to synapse dynamics and centralization. Measurement of the distance across T cell microclusters following TIRF imaging of synapses – a means of assessing centralization – revealed that dimeric CAR microclusters only formed a central dense cluster under conditions of low affinity and low antigen density, whereas the monomers reliably formed this structure (**Fig. 5a-b**).

**Figure 5:**
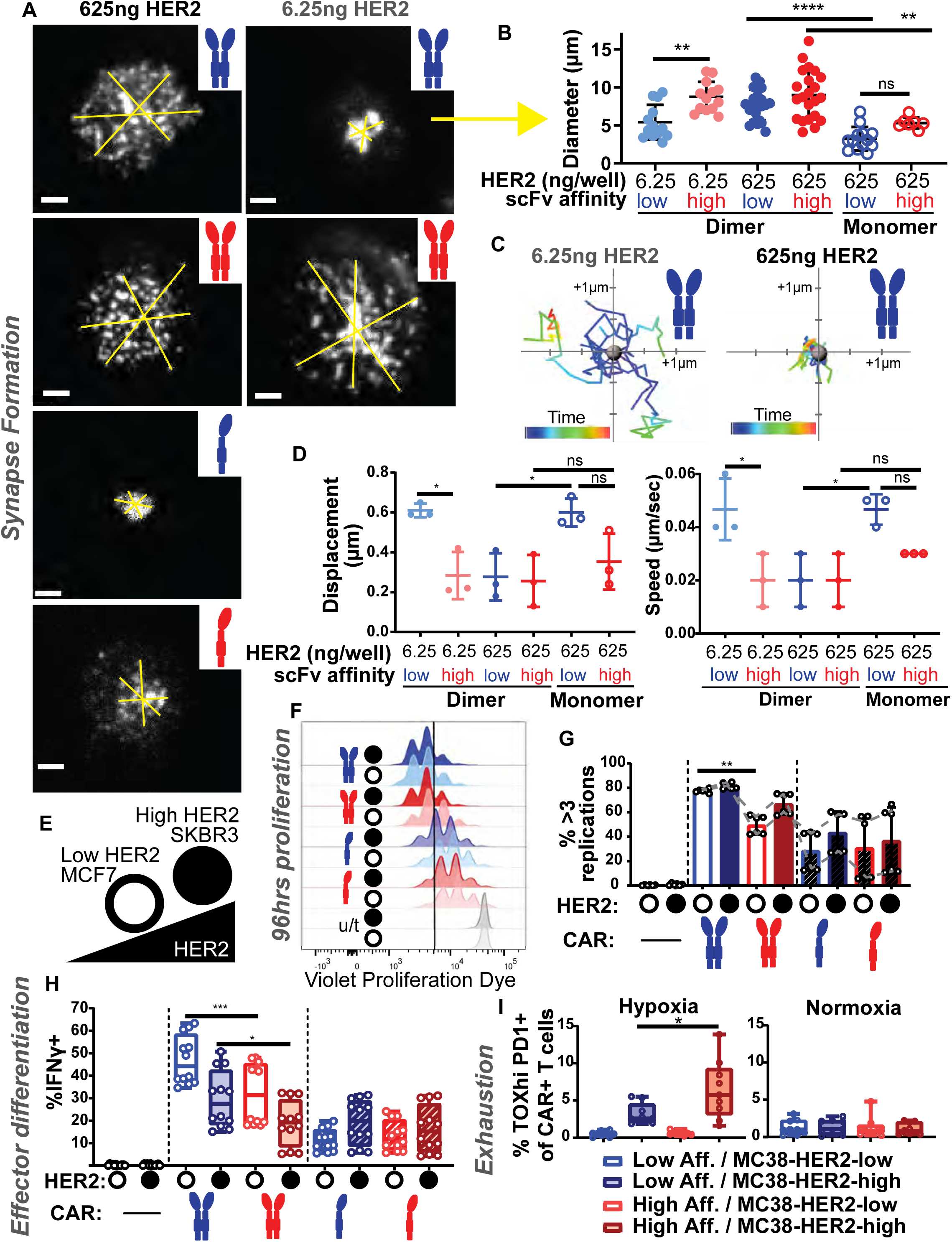
Impaired movement of high affinity dimer CAR microclusters in synapses and reduced effector function. A-B. TIRF imaging of CAR-mEmerald is shown for synapses with low- or high-affinity CAR interacting with low or high HER2-loaded bilayers (A). Time point shown is 93.6 seconds following the initiation of imaging, which began as synapses were starting to form. Yellow lines were drawn to span CAR microclusters in synapse, indicating diameter quantified in B. Three lines were averaged and line assignment was blinded to account for manual drawing. For dimeric CARs, only low affinity:low HER2 synapses result in accumulation of CAR microclusters at the center, indicated by lower diameter. Monomerization of CARs improves centralization on high HER2. Data is shown for at least 7 cells pooled from a minimum two independent experiments per condition (n = 13, 13, 20, 20, 13, 7 cells per group from left to right, respectively). Error bars represent s.d. Analyses shown are Šídák’s multiple comparisons tests. Scale bars = 2 μm. C-D. Spots with tracking were assigned to CAR microclusters in Imaris. Examples of mobile (left, low affinity dimer on low HER2) and immobile (right, low affinity dimer on high HER2) microclusters are shown as a flower plot for 10 random tracks (C). Quantification is shown for the average displacement (left) and speed (right) of all CAR microclusters for a given cell (D, n = 3 cells per group). Limited mobility is apparent for dimeric CARs on high-HER2 loaded bilayers. On low-HER2 bilayers, low-affinity CAR microclusters show increased mobility. Error bars represent s.d. Analyses shown are Šídák’s multiple comparisons tests. E. Low HER2-expressing MCF7 (open circle) and high HER2-expressing SKBR3 (solid circle) cell lines were used to assess differences between HER2 levels in cell-cell interactions in vitro. F. Proliferation is induced across all CAR+ conditions with low and high HER2, as seen by VPD dilution at 96 hrs following co-incubation. Line marks VPD dilution indicating at least 3 replication cycles (quantified in F). G. Percentage of cells that have undergone at least 3 replication cycles. Replicates from 2 independent experiments of different donors are pooled (n = 6). Dashed lines indicate individual donor trends. Error bars represent s.d. Analysis shown is Šídák’s multiple comparisons test. H. Intracellular staining with anti-IFN-γ-APC at 18hrs following co-incubation. Low-affinity CAR induces greater IFN-γ than high-affinity. Monomeric low-affinity CAR has overall lower magnitude of response, but improves the dose response. Box and whiskers plot error bars showing minimum and maximum values (n = 12 samples per group pooled from four independent experiments and two donors). Analyses shown are Šídák’s multiple comparisons tests. I. Low-affinity (blue) and high-affinity (red) CARs were retrovirally expressed in primary mouse CD8+ T cells and co-cultured with MC38s expressing low- or high-HER2 for 24 hrs in normoxia (20% oxygen) before separation into normoxic or hypoxic (1.5% oxygen) cultures for an additional six days. Hypoxic co-cultures of high-affinity CAR T cells with MC38-HER2-high cells have significantly more TOX high/ PD1+ cells than low-affinity CAR T cells, indicative of development towards an exhausted state. Box and whiskers plot error bars showing minimum and maximum values (n = 9 samples pooled from 3 independent experiments). Analysis shown is unpaired t test.

In order to determine whether this phenotype was the result of a lack of mobility of microclusters, as opposed to direct accumulation of CARs in the center vs. periphery, we tracked the microclusters over time (**Supplemental Fig. 3a** and **Supplemental Videos 5-10**). On high-HER2 bilayers, the low-affinity CAR monomer yielded mobile microclusters, whereas both CAR dimers showed impaired mobility as demonstrated by flower plots of microcluster displacement, and analysis of displacement and speed (**Fig. 5c-d** and **Supplemental Fig. 3b-c**). The mobility of low-affinity but not high-affinity CAR dimer microclusters was restored on low HER2-loaded bilayers (**Fig. 5c-d** and **Supplemental Fig. 3b-c**). Analysis of the direction of microcluster movement revealed no significant differences (**Supplemental Fig. 3d**). Therefore, we found that a monomerized low-affinity CAR supported the formation of synapse architectures and dynamics characteristic of TCRs on antigen-loaded lipid bilayers, whereas high-affinity dimerized CARs did not.

Given these proximal effects on synapse dynamics, we sought to determine the downstream effects of the hyper-stable or more physiological receptors. In order to compare between HER2 levels for T cell stimulation experiments, we first used two breast cancer cell lines: the MCF7 luminal breast cancer cell line, which does not over-express HER2 (Nobili et al., 2021; Comsa et al., 2015; Liu et al., 2015), and the SKBR3 cell line, which endogenously over-expresses HER2 and thus mirrors HER2-positive breast cancers (Mota et al., 2017) (**Fig. 5e** and **Supplemental Fig. 2h, 3e**). In coupling assays to assess overall synapse formation (Friedman et al., 2006), all combinations yielded robust coupling (**Supplemental Fig. 3f-i**). Early activation, as measured by CD69 staining at 18 hours following stimulation, and entry into cell cycle, measured by Ki67 staining at 96 hours, were similarly robust for all CAR interactions with either low- or high-HER2 targets (**Supplemental Fig. 3j-l**). While all conditions induced a proliferative burst as measured by VPD dilution after 96 hours, rounds of proliferation varied significantly, with the greatest proliferation observed using low-affinity dimeric CAR (**Fig. 5f-g**).

Assaying effector function by intracellular staining for IFN-γ at 18 hours post-stimulation also revealed increased production of this cytokine in cells bearing low-affinity CAR dimers relative to high-affinity ones (**Fig. 5h**). In these cocultures, less IFN-γ was produced by dimeric CAR T cells when incubated with high-HER2 SKBR3s than with low-HER2 MCF7s. In this regard, while the low-affinity monomeric CAR displayed a decreased magnitude of response, likely in part due to a lack of constitutively paired intracellular CD3ζ, it improved the expected dose-dependent response pattern.

Finally, we incubated mouse CAR T cells with MC38 tumor cells expressing variable HER2 levels – using this strategy to also minimize the effect of varying the background gene effects of using two cell lines. Low and high affinity receptors performed similarly for cytokine production (**Supplementary Fig. 3m-n**) but when we shifted these cultures to hypoxia, mimicking the tumor microenvironment and licensing deeper progression into exhaustion (Scharping et al., 2021), the conventional high affinity dimer was more susceptible to exhaustion as measured by TOX and PD1 co-expression (**Fig. 5i**), in line with recent work (Shakiba et al., 2021) using native TCRs.

## Discussion

Factors that control the dynamics of T cell microvilli are not well understood. This is in part because this dynamic microvillar biology is somewhat unique – the well-studied microvilli at the intestinal brush border are relatively much less dynamic (Crawley et al., 2014; Meenderink et al., 2019). Here, using LLS and synaptic contact mapping TIRF imaging, we show that CARs and TCRs distribute as patches that occupy regions of both microvilli and cell cortex on isolated T cells much like TCR, although not obligately overlapping in position. The distinctness of the synthetically engineered transmembrane domain – taken from CD8 – and intracellular domains appear to result in an initial propensity to segregate as patches independently of TCRs. We hypothesize that this results from differential miscibility and recruitment of membrane lipids related to combinations of transmembrane and cytoplasmic domains although studies of this represent a significant tangent from our work. Future studies will be needed to reveal the factors that lead to the formation of patches, or “condensates” (Lyon et al., 2021), prior to TCR triggering and how to manipulate them. Following receptor binding, we show that CAR microclusters accumulate at the tips of microvilli, which are hyperstabilized in comparison to those mediated by TCRs. The finding that contact stability is directly related to receptor affinity and avidity is consistent with previous work showing that stabilization by TCR was maintained even following inhibition of TCR-proximal signaling or actin blockade (Cai et al., 2017). Taken together, this suggests that microvillar stabilization is highly related to the strength of physical binding of extracellular receptors.

The actin cytoskeleton is a key regulator of T cell function, not just for successful migration (Thompson et al., 2021), but also for optimal immune synapse formation and T cell activation (Dupré et al., 2021). In fact, even the T cell’s synaptic partners can contribute to the cytoskeletal dance – dendritic cell (DC) actin dynamics promote efficient T cell activation, and notably favor a multifocal distribution of microclusters at the T:DC synapse (Leithner et al., 2021). A multifocal synapse also forms during thymic negative selection (Richie et al., 2002), where TCRs that bind too strongly to self-ligands induce cell death. Slower receptor unbinding kinetics have also been associated with reduced actin velocity (Colin-York et al., 2019). Given these findings, it is tempting to speculate that multifocal patterns may be a general indicator of relatively “strong” contacts (Mossman et al., 2005), although as the above examples demonstrate, the implications of that phenotype vary by context. We see here on SLBs that it represents a super-physiological outcome of ultra-high affinity/avidity binding that does not correlate with optimal effector function. Many prior works also demonstrate that increasing the binding strength of antigen receptors does not reliably predict the efficiency of T cell activation and ultimate effector function, both for natural TCR (Corse et al., 2010; McMahan et al., 2006; Kalergis et al., 2001; Schmid et al., 2010; Dushek et al., 2011; Hebeisen et al., 2013) and engineered CAR (Harris et al., 2018; Ghorashian et al., 2019; Park et al., 2017; Liu et al., 2015).

Future studies to further characterize the regulators of T cell microvillar dynamics and better sub- classify actin-based synaptic protrusions, especially those present during early and late synapses, will reveal new mechanisms of control of T cell activation (Blumenthal and Burkhardt, 2020). The experiments performed here reveal that receptor binding affinity and avidity control the dynamics of synaptic microvilli. Given that microvillar persistence in TCR-occupied contacts was not dependent on downstream signaling through ZAP70 (Cai et al., 2017), these findings indicate that affinity and avidity of binding directly affect the movements of the opposing cell surfaces. Thus, discovering how engineered receptors interact with the cell biological processes underlying recognition contributes to our understanding of the mechanism both of the therapy and the system itself. This work opens up new avenues for future research to determine whether better quality T cells could be produced for cell therapies by boosting the intracellular signal downstream of physiologically normalized binding dynamics. The formation of receptor-ligand complexes at cell-cell interfaces is integral to intercellular communication. However, the 3-dimensional structure of cellular interfaces and how movements of the plasma membrane affect the formation of and signaling through such complexes is relatively understudied. Similar mechanisms are likely at play in other cell:cell interfaces.

## Author Contributions

C.B. designed, conducted, and analyzed most experiments, and drafted the manuscript. J.E. and K.M. authored unique MatLab codes and discussed data. K.M. also assisted with LLS imaging acquisition. E.C. discussed project design. C.A.C. assisted with experiments and discussed data.

V.S. designed affinity measurement experiments. K.T.R. contributed to the formulation of the project, designed experiments, provided reagents for experiments involving human T cells, and interpreted data. M.F.K. contributed to the formulation of the project, designed experiments, interpreted data, and developed the manuscript. All authors contributed manuscript revisions.

## Data and Materials Availability

The datasets and unique materials generated during the current study are available from the corresponding author on reasonable request.

## Code Availability

Unique analysis codes have been made available and can be accessed via the GitHub links provided in the relevant methods sections.

## Materials and Methods

### Lentiviral and retroviral CAR constructs

All CARs were fused to C-terminal MYC tag, CD8α hinge/transmembrane domain, 4-1BB co-stimulatory domain, CD3ζ signaling domain, and an N-terminal mEmerald tag. Monomeric versions of each CAR were created by using the Q5 Site Directed Mutagenesis Kit (NEB #E0554S) yielding two cysteine to serine point mutations in the CD8α hinge (Hennecke and Cosson, 1993) (**Supplemental Table 1**).

### Human T cell culture, lentiviral transduction, and co-incubations

Lenti-X 293T cells (Takara Bio) were transfected with pHR SIN including cloned transgene and packaging vectors pMD2.G and pCMVdR8.91 using TransIT-Lenti Transfection Reagent (Mirus #MIR6603). On the day of transfection, primary human CD8+ T cells were thawed into complete human T cell media: X-VIVO 15 (Lonza # 04-418Q), 5% Human AB serum (Valley Biomedical #HP1022) and 10 mM neutralized N-acetyl L-Cysteine (Sigma Aldrich #A9165-25G) supplemented with 30 U/mL recombinant human IL-2 (R&D Systems #202-IL), and 55 μM beta-mercaptoethanol (Thermo # 21985023). The following day, Human T-Activator CD3/CD28 Dynabeads (Thermo #11161D) were added at 1:1 ratio with thawed T cells. The next day, T cell media was replaced with Lenti-X 293T viral supernatant. For lentiviral transduction of monomeric CARs, virus was concentrated by PEG/NaCl precipitation, and stored at -80° C prior to use. Viral supernatant was replaced with fresh T cell media the next day, and T cells were allowed to recover for one day prior to Dynabead removal and sort. Cells were sorted for CAR-mEmerald expression in the range of 1-2 logs above background. Cells were then rested and used at 10-21 days post initial stimulation. Lenti-X 293T cells were cultured in Dulbecco’s modified Eagle’s medium (DMEM, Gibco #11995) with 10% fetal bovine serum (MilliporeSigma), penicillin (50 U/ml) and streptomycin (50 μg/ml) (MP Biochemicals #MP091670049), and 1 mM sodium pyruvate (Sigma-Aldrich #S8636). For early activation and intracellular cytokine assays, 5x10^4^ T cells were added at a 1:1 ratio to 96-well flat-bottom plates with MCF7 (ATCC) or SKBR3 (ATCC) cells for 18 hours. BD GolgiPlug (#555029) was added for the final 10 hours. For proliferation assays, T cells were stained with Violet Proliferation Dye (BD #562158) prior to plating of 2x10^4^ T cells at 1:1 ratio with MCF7s or SKBR3s. Complete T cell media was supplemented the following day, and cells were analyzed by flow cytometry at 96 hours following plating.

### Flow cytometric analysis

Zombie NIR Fixable Viability Kit (BioLegend #423106) was used for exclusion of dead cells. Surface staining was performed with anti-mouse Fc receptor antibody (clone 2.4G2, UCSF Hybridoma Core) or Human TruStain FcX (BioLegend #422302) in PBS with 2% FCS for 30 min on ice. Supplemental Table 2 lists all antibodies referenced for flow cytometry and imaging experiments. For experiments with staining of nuclear proteins, eBioscience Foxp3 / Transcription Factor Staining Buffer Set (Fisher Scientific #00-5523-00) was used for fixation and permeabilization. For all other experiments involving intracellular staining, BD Cytofix/Cytoperm (#554722) was used. Following fixation and permeabilization, cells were incubated with Fc block for 10 min on ice prior to addition of intracellular stain. Flow cytometry was performed on a BD Fortessa instrument, and sorting was performed on BD FACSAria or BD FACSAria Fusion instruments. FlowJo software (BD Biosciences) was used for all analyses. Flow-cytometry based coupling assay was performed as previously described(Friedman et al., 2006) on unfixed cells using DAPI for live/dead discrimination.

### Mice

C57BL/6J and B6.SJL-Ptprc^a^ Pepc^b^/BoyJ (CD45.1) mice, used as sources of primary mouse T cells, were housed and bred at the University of California, San Francisco, according to Laboratory Animal Resource Center guidelines. Protocols were approved by the Institutional Animal Care and Use Committee of the University of California.

### MC38-HER2 retroviral transduction

Truncated HER2, without intracellular signaling domains, (NP_004439.2; amino acids 1 to 730) was cloned into pIB2 retroviral vector. Phoenix cells were transfected using FuGENE 6 Transfection Reagent (Promega #E2691), and retroviral supernatant was collected and used immediately for transfection of MC38s on days 2 and 3 following transfection. Two days after the second transduction, cells were sorted for expression of HER2 (stained with anti-HER2-A488). MC38-HER2 cells were then expanded and the retroviral transduction process was repeated. Following the second transduction, cells were then sorted into high-, medium-, and low-expression levels using MCF7 and SKBR3 cells as standards for low and high expression, respectively. MC38 cells were cultured in DMEM (Gibco #11995) supplemented with 10% fetal bovine serum (Benchmark), 100 U/mL penicillin, 0.1 mg/mL streptomycin, 2 mM L-glutamine (Gibco #10378), 10 mM HEPES (Thermo #15630106), and 55 μM beta-mercaptoethanol (Thermo # 21985023).

### Murine T cell culture, retroviral transduction, and functional assays

The mouse OT-I TCR system was chosen as a comparator to the anti-HER2 CAR for its affinity near the top of the common range, and in order to avoid double-transfection (of a human TCR along with a CAR) which otherwise would create significant experimental inefficiencies. For all experiments using murine T cells, cells were maintained in RPMI (Gibco #11875) supplemented with 10% fetal bovine serum (Benchmark), 100 U/mL penicillin, 0.1 mg/mL streptomycin, 2 mM L-glutamine, 10 mM HEPES (Thermo #15630106), 55 μM beta-mercaptoethanol (Thermo # 21985023), non-essential amino acids (Thermo #11140050), 1mM sodium pyruvate (Sigma-Aldrich #S8636), and supplemented with 100 U/ml IL-2, which is referred to as complete RPMI. Single cell suspensions were prepared from the lymph nodes and spleens of C57BL/6J, *Ptprc^a^* (CD45.1), or OT-I TCR transgenic mice. Following red blood cell lysis of splenocytes, negative selection using the EasySep Mouse T Cell or CD8+ T cell Isolation Kit (STEMCELL Technologies #19853) was used to isolate CD8+ T cells. T cells were activated in complete RPMI using CD3/CD28 Mouse T activator Dynabeads (Thermo #11-453-D) for 24 hours before the first round of retroviral transduction.

For retrovirus production, Platinum-E cells were transfected with pMIG including CAR transgene using FuGene. Transfections were performed in DMEM (Gibco #11995) supplemented with 10% fetal bovine serum (Benchmark) and 10 mM HEPES (Thermo #15630106), which was replaced with complete RPMI (without IL-2) the following day. Retroviral supernatants were harvested at day 2 and 3 and stored in -80°C. Platinum-E cells were maintained in DMEM (Gibco #11995) supplemented with 10% fetal bovine serum (Benchmark), 100 U/mL penicillin, 0.1 mg/mL streptomycin, 2 mM L-glutamine (Gibco #10378), 10 mM HEPES (Thermo #15630106), 10 µg/ml blasticidin (Fisher Scientific #A1113903) and 1 µg/ml puromycin (Gibco #A1113803).

For T cell retroviral transduction, retroviral supernatant was added to T cells in retronectin-coated plates at 24 and 48 hours following initial stimulation, and the plates were centrifuged for 1 hour at 2,000g and 30°C. After the second spinfection, cells were rested two days prior to Dynabead removal (4 days post-stim). T cells were then sorted for imaging or rested for an additional 6-7 days in 10 ng/ml recombinant murine IL-7 (PeproTech #217-17) and 100 U/ml IL-2 and used for in vitro co-culture experiments.

For binned HER2 expression level experiments, MC38-HER2 lines were sorted into 5 consecutive bins by HER2 expression using MCF7s and SKBR3s as the low and high HER2 standards, respectively. MC38-HER2 cells, sorted as described, were then plated at 5x10^4^ cells/well in flat-bottom 96 well plates, and 5x10^4^ T cells were then added, bringing the total volume to 200 µl/well complete RPMI. For hypoxia experiments, T cells were rested 6 days and then plated at 5x10^4^ cells at a 1:1 ratio with MC38-HER2-high or MC38-HER2-low cells in two 96-well plates in complete RPMI. At 24 hours, wells were replenished with complete RPMI + IL-2, and 5x10^4^ MC38- HER2-high and -low cells were added. One plate was moved to 1.5% oxygen while the second was maintained at 20% oxygen. MC38-HER2-high and -low cells and media + IL-2 were then replenished every two days until analysis on day 7 after start of co-culture (6 days in hypoxia).

### Surface plasmon resonance affinity measurements

Measurements were taken using a Biacore T200 instrument with CM4 sensor chip and HBs-EP+ buffer. HER2-mIgG2aFc (ACROBiosystems #HE2-H5255) was captured using anti-mIgG (50 RUs). Association and dissociation times were 120 and 900 sec, respectively. Concentrations of scFv used for single cycle kinetics analysis: 0.33, 1, 3, 9, 27 nM.

### Lattice Light-Sheet Microscopy

Lattice light-sheet imaging was performed as described previously(Cai et al., 2017). 5 mm round coverslips were cleaned by a plasma cleaner and coated with 2 μg/ml fibronectin in PBS at 37°C for 1 hour, or at 4°C overnight, before use. ∼3×10^5^ CAR T cells were loaded onto the coverslip and incubated at 37°C for 30 min. Cells were then fixed in d_2_h_2_0 with 20 mM HEPES (Thermo #15630106), 0.2 M sucrose (RPI #S240600, 4% paraformaldehyde (Election Microscopy Sciences #15710), and 8% glut-aldehyde (Election Microscopy Sciences #16019) for 10 min at room temperature. Coverslip was washed gently in 1 ml PBS and then stained with antibodies to CD45 and/or MYC with anti-mouse Fc receptor antibody (clone 2.4G2, UCSF Hybridoma Core). Samples were stained for at least 30 min and kept at 4°C until use. Prior to imaging, coverslip was gently washed with 1 ml warmed RPMI without phenol red (Gibco #11835) supplemented with 2% fetal bovine serum, 100 U/mL penicillin, 0.1 mg/mL streptomycin, 2 mM L-glutamine, 10 mM HEPES and 50 µM β-mercaptoethanol (imaging media). For imaging of live CAR T cell interactions with MCF7 and SKBR3 targets, MCF7 or SKBR3 cells were plated onto fibronectin-coated coverslips 1-2 days prior to imaging, or onto Cell-Tak (Corning #354240) coated coverslips with a 10 min spin at 1400 rpm and 4°C. CAR T cells were stained with antibody to CD45- Alexa647 for 30 min on ice. Target cells on coverslip were stained with CFSE (Invitrogen #C34554) for 20 min at 37°C, or were identified by nuclear-localized mKate expression. Cells were then washed and T cells were added onto the coverslip prior to being loaded into the sample bath with warmed imaging media and secured. Imaging was performed with a 488 nm, 560 nm, or 642 nm laser (MPBC, Canada) dependent upon sample labeling. Exposure time was 10 ms per frame leading to a temporal resolution of ∼4.5 and ∼6.75 seconds in two- and three-color mode respectively.

### Supported lipid bilayers, synaptic contact mapping, and calcium flux imaging

Preparation and use of supported lipid bilayers was performed as described previously (Cai et al., 2017; Beemiller et al., 2012b). Mixtures of 96.5% POPC, 2% DGS-NTA (Ni), 1% Biotinyl-Cap-PE and 0.5% PEG5,000-PE (Avanti Polar Lipids #850457C, 790404C, 870273C, 880230C) were made in a round bottom flask and dried under a stream of nitrogen and then overnight under vacuum. The phospholipids were then rehydrated at a total concentration of 4 mM in PBS for one hour to create crude liposomes. Small, unilamellar liposomes were then made by extruding through 100 nm Track Etch filter papers (Whatman #800309) with an Avestin LiposoFast Extruder (Avestin).

8-well Nunc Lab-Tek II chambered coverglass (Thermo #155360) were cleaned by submersion in 5% Hellmanex III (Sigma-Aldrich #Z805939). Flask containing chamber in solution was microwaved for 25 sec and then allowed to clean at room temperature overnight. The chambers were then washed repeatedly with 18 Milli-Q water and then dried. Finally, 250 µl 3M NaOH was added to each well for 15 min at 55°C. Wells were washed with 300 µl Milli-Q water and the NaOH cleaning was repeated. Wells were then washed thoroughly and dried prior to use. Lipid bilayers were set up on the chambered coverglass by adding 0.25 mL of a 0.4 mM liposome solution to the wells. After 30 minutes, wells were rinsed with 8 mL of PBS by repeated addition of 0.5 mL of PBS, then aspiration of 0.5 mL of the overlay. Non-specific binding sites were then blocked with 1% BSA in PBS for 30 minutes. After blocking, 25 ng of unlabeled streptavidin (Invitrogen #43- 4301) was added to each well and allowed to bind to bilayers for 30 minutes. After rinsing, protein mixes containing 63 ng recombinant human ICAM-1 (Abcam #AB151393) and 6.25-625 ng biotinylated HER2 (ACROBiosystems #HE2-H822R) or 6 ng pMHC in 2% BSA were injected into each well. pMHC was provided by the NIH Tetramer Facility. After binding for 30 minutes, wells were rinsed again and 25 ng of QDot605-streptavidin (Thermo #Q10101MP) was added to each well. For calcium flux imaging, QDot605-streptavidin was not added. Bilayers were finally rinsed with imaging media before being heated to 37°C for experiments. Experiments using 5 µm diameter silica microspheres (Bangs Laboratories #SS05003/SS05N) were performed as previously described (Beemiller et al., 2012a). Briefly, the same protocol was used for building a lipid bilayer on chamber coverglass as for 4x10^5^ beads, based off equivalent surface area, but with washes performed by centrifugation instead of repeated aspiration.

For synaptic contact mapping (SCM) experiments, 5×10^5^ T cells were added to the well prior to imaging. Once cells began interacting with the bilayer, imaging was initiated. For imaging of OT-I TCR, 1-2×10^6^ OT-I T cells were stained with 2.5 µg H57-597 non-blocking monoclonal antibody conjugated to Alexa Fluor 488 on ice for 30 minutes, then rinsed once with complete imaging media. Imaging for synaptic contact mapping was performed as described previously(Cai et al., 2017). The TIRF microscope is based on a Zeiss Axiovert 200M equipped with a 100x 1.45NA oil immersion objective, DG-4 Xenon light source (Sutter) and Zeiss TIRF slider (Cai et al., 2017; Beemiller et al., 2012b). All images were collected using a DV2 image splitter (Photometrics) positioned in front of an Evolve EMCCD (Photometrics). A 4 band multi-color TIRF dichroic located in the microscope separated the excitation and emission light for imaging (Chroma Technology). Images of CAR or TCR were collected by imaging CAR-mEmerald (or Alexa Fluor 488-labeled TCRs) using TIRF mode, by imaging QD605-streptavidin in widefield mode, and by imaging cells with interference reflection microscopy (IRM), also in widefield mode. Widefield QD605-strepavidin images were collected using a 405/10 nm excitation filter (Chroma Technology) located in the DG4 light source, while samples imaged with TIRF were excited by an Obis 488nm laser (Coherent). IRM images were acquired using a 635/20 nm excitation filter (Chroma Technology) positioned in the DG4 light source. The CAR/TCR and QD605 emitted fluorescence signals were separated using a DV2 image splitter with a 565 nm long-pass dichroic mirror installed along with 520/35 nm and 605/70 nm emission filters (Chroma). Images containing the IRM signal were acquired through the long-pass dichroic and 605/70 nm emission filter in the image splitter. For Fura-2 imaging, cells were stained with 2 µM Fura-2 dye (Thermo #F1221) for 15 min at room temperature. Cells were then washed in imaging media, and 5×10^5^ T cells were added to the imaging well. 3 min after addition of cells, acquisition was initiated. Fura-2 imaging experiments were acquired using a 40x 1.3NA oil immersion objective (Zeiss) and the same light source and dichroic described above. Widefield Fura-2 340 nm and 380 nm images were collected using 340/26 (Semrock, Rochester, NY) and 380/30 (Chroma, VT) excitation filters, respectively, located in the DG4 light source.

### Image analysis

All computational image analysis for SCM imaging was performed in Matlab (The Mathworks, Natick, MA), Imaris version 9.2.1 or 7.6.3 (Bitplane), and Fiji. CAR-mEmerald microcluster tracking analysis was performed using the spots function in Imaris with the following parameters:

0.25 µm estimated diameter, autoregressive motion tracking, 0.5 µm maximum distance, gap size 3, track duration > 5 sec. Centroid positions for dot product calculations were defined by making a surface of the synapse CAR-mEmerald interface in Imaris with grain size of 3 µm and largest sphere diameter of 1 µm. Analysis for LLS was performed in Imaris and Matlab. Unique analysis code has been made available through GitHub and can be found at the URLs included below.

### Lattice Light-Sheet: post processing

Raw data were deconvolved using the iterative Richardson-Lucy deconvolution process with a known point spread function that was recorded for each color prior to the experiment, as described previously (Cai et al., 2017). A typical sample area underwent 15-20 iterations of deconvolution. For live imaging experiments, photobleaching correction was applied in FIJI using the histogram matching method.

### Close contact segmentation and persistence analysis

Close contact segmentation, CAR co-localization, and persistence time analysis was performed using Matlab and Imaris as previously described (Cai et al., 2017). Briefly, the IRM images were used to define the region of the cell interface. Active contour segmentation of the QD605 image was then used to define close contact regions. These regions were then converted to Spots objects in Imaris. CAR intensity was masked to regions of close contacts, and the average intensity for each contact area was then plotted in a histogram. A Gaussian distribution curve centered at the background fluorescence median was overlayed. Contacts that fell within 3 sigma of the Gaussian distribution were considered CAR^-^, while the higher intensity contacts were considered CAR^+^. These contacts were then separated into separate image stacks and persistence time for individual contacts were calculated. Contact persistence time was determined by summing the number of frames each binary connected component object existed for and multiplying by the time per frame. Contacts were assumed to not travel more than their diameter per time point. Code for analysis of persistence times has been made available: https://github.com/BIDCatUCSF/NanocontactsTIRF_V5.

### Radial intensity profiles

Definition of cell boundary, assignment of radial coordinates, and plotting of pixel intensities were performed using MatLab (Natick, MA). The outer edge (boundary) of cells was detected using a custom program which primarily applied a two-step kmeans clustering calculation on each of the image slices collected in the z-stack of images describing a single cell. The boundary was then eroded by three pixels to accommodate the resolution of the LLS imaging system. Code for defining the cell boundary: https://github.com/BIDCatUCSF/Exterior-t-Cell-Edge-Detection. Radial coordinates were assigned for plotting of radial intensity profiles using this code: https://github.com/BIDCatUCSF/Outer-Boundary-Profile-Code. Binned intensity was measured by taking a moving average of ten pixels. Excursions of the binned intensity above the channel mean intensity + 3 s.d. were used as the threshold to define patches.

### Three-dimensional surface curvature mapping and patch analysis

The surface curvature and patch analysis used here was performed in MatLab as described in Cai et al. 2022 (Cai et al., 2022). Briefly, the cell boundaries (as defined above for radial intensity profiles) were used to calculate Pearson’s correlation coefficients for variable sized kernels in three dimensions, roughly of the shape of the point spread function. These correlation coefficients were then used as the basis for clustering analysis by segmentation and watershed, creating the patches. Surface curvature was calculated for each position of the cell boundary, mapped by color, and thresholded to regions of low curvature to indicate peaks on the cell surface. Projections of the surface curvature and receptor intensity onto two dimensions were created using Map3-2D software (Sendra et al., 2015).

### Fura-2 ratio image analysis

Fura-2 340nm/380nm ratio images were created in MetaMorph Version 7.6.5.0 using a maximum ratio of 7 and imported to Imaris for tracking using the surface function with the following parameters: 1 µm grain size, 0.75 µm diameter of largest sphere, 2 µm region growing estimated diameter autoregressive motion tracking, 2 µm maximum distance, gap size 1, track duration above 148 sec. For analysis of final time point, surfaces were made for all cells in field of view without tracking: 0.75 µm grain size, 0.75 µm diameter of largest sphere, 1.5 µm region growing estimated diameter. Maximum 340nm/380nm ratio per cell was compared for all cells in field of view at final imaging time point – 5.5 min after addition of T cells to chamber well.

### Statistics

Statistical tests were performed using GraphPad Prism Version 9.0.1. Independent experiments and donors are as noted in figure legends – all other replicates are technical replicates. Significance tests are as described in legends and include: Tukey’s multiple comparisons tests, Šídák’s multiple comparisons tests, and two-tailed t tests. Scatter plots show mean and s.d (error bars). Box and whisker plots indicate median, 25^th^ to 75^th^ percentile (box), and minimum to maximum (whiskers). P values are reported as follows: ≥ 0.05 as ns, 0.01 to 0.05 as *, 0.001 to 0.01 as **, 0.0001 to 0.001 as ***, and < 0.0001 as ****.

## Supplemental Figure Legends

**Supplemental Figure 1.**
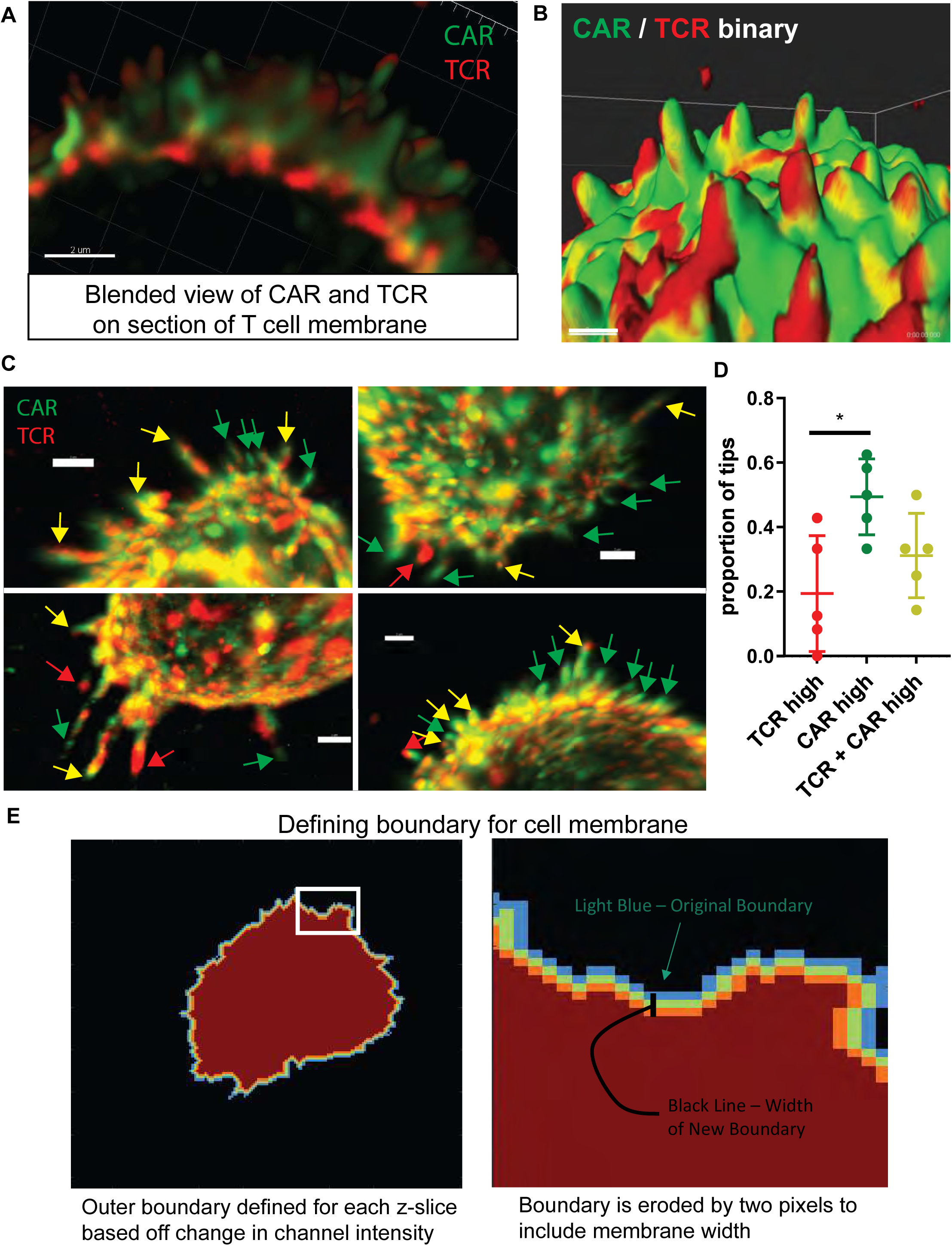
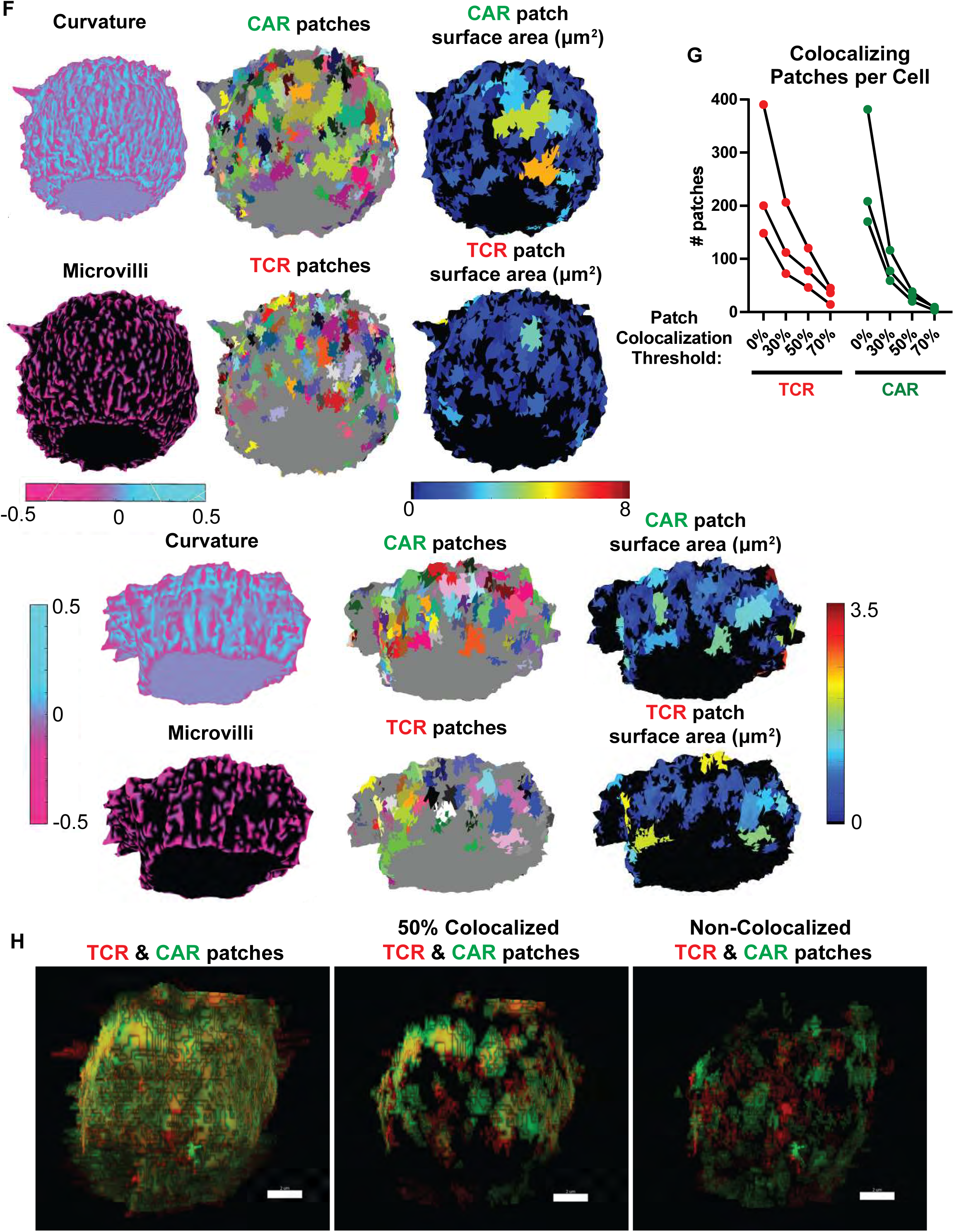
Imaging TCR and CAR on the surface of isolated T cells. A-B. Blended view (A) and shaded binary (B) Imaris rendering of fixed anti-HER2 CAR T cell surface, as described in Fig. 1A. Blended rendering shows section of 40 z slices. Scale bars are 2 μm. C. Regions of interest (in addition to Fig. 1B) that were used for the manual count of microvillar tip receptor occupancy in Fig. 1C and (D). Microvilli tips are marked as being high for TCR (red arrow), CAR (green arrow), or both (yellow arrow). Scale bars = 2 μm. D. Proportion of microvillar tips per region of interest that were marked as high for, CAR, or both. 41 tips across 5 regions and 3 cells were scored. E. Schematic of method used to define the cell membrane boundary used in radial intensity profiles. F. Two additional examples of cells analyzed as described in Figure 2A-F. G. Clusters are quantified for increasing thresholds of co-localization. Left: TCR clusters are quantified which have 0, 30, 50, or 70% overlap with CAR clusters. Right: CAR clusters are quantified which have 0, 30, 50, or 70% overlap with TCR clusters. H. Maximum intensity projection of TCR and CAR patches rendered in Imaris. For each patch, the intensity is defined by the integrated intensity for that patch. All patches are shown on the left. Patches which are ≥50% co-localized (i.e. 50% of the CAR patch overlaps with TCR, and vice versa) and <50% co-localized are shown in middle and left panels, respectively.

**Supplemental Figure 2.**
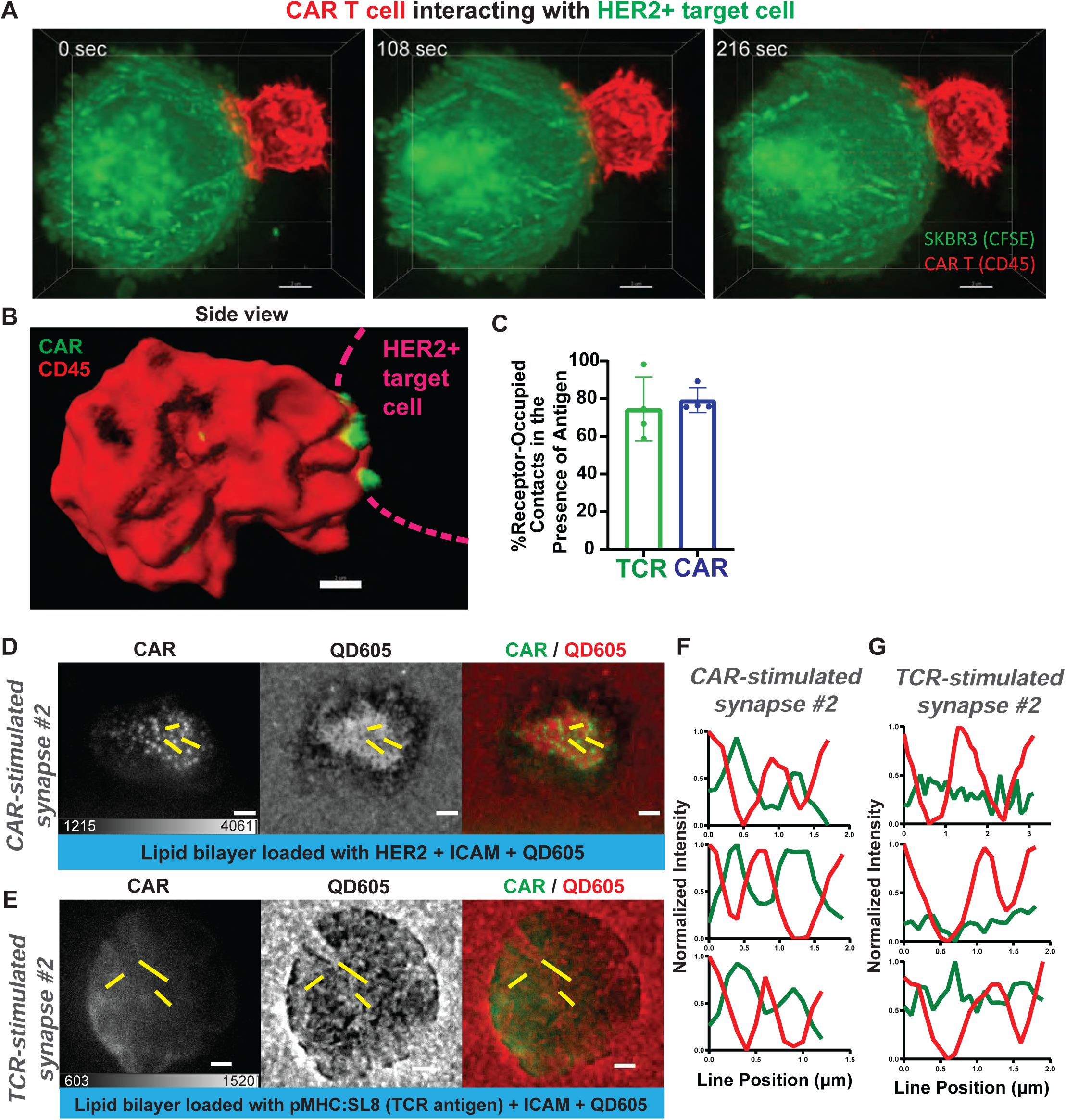
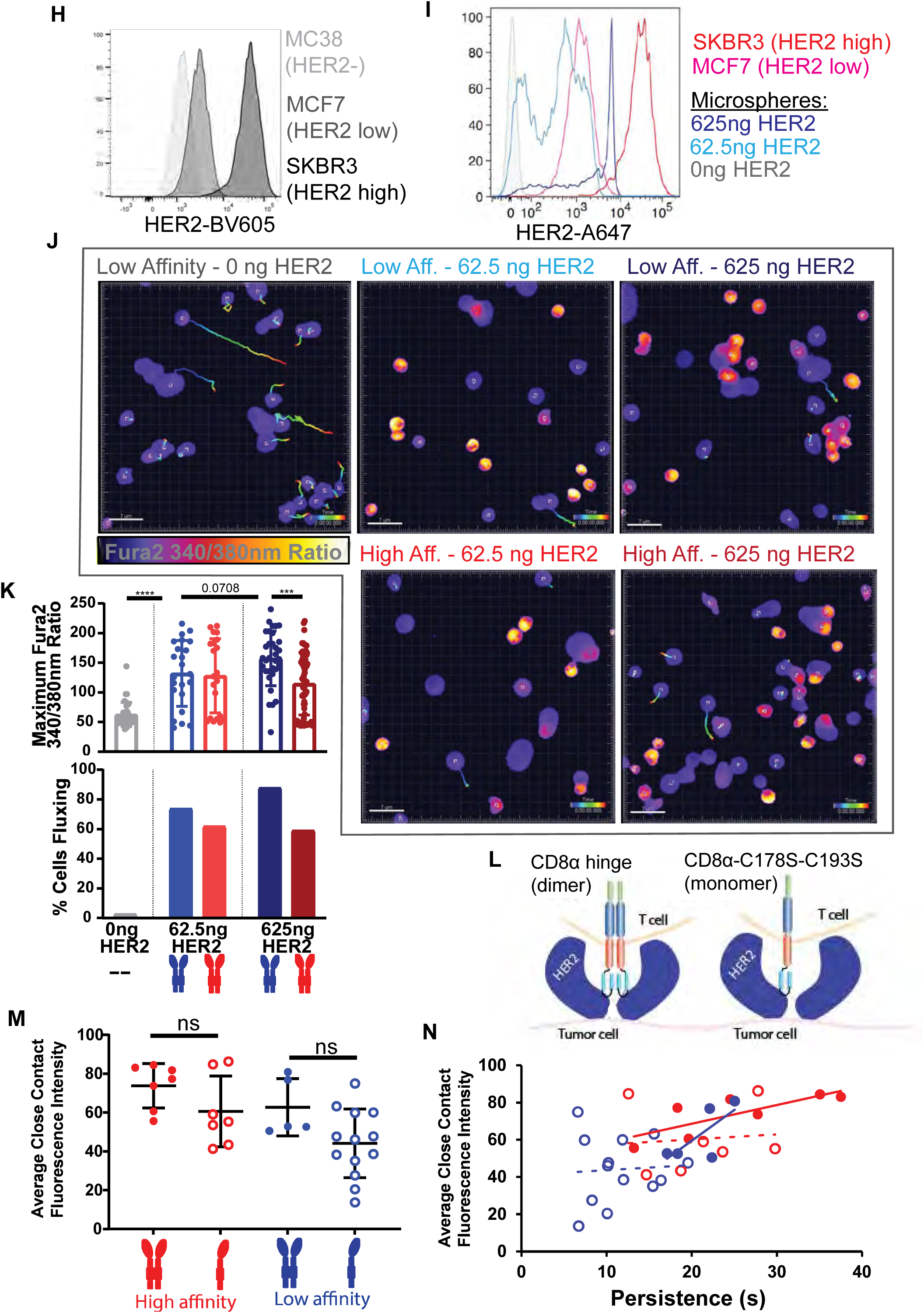
Imaging CAR T cell:target cell interactions across affinity and antigen densities. A. Time-lapse of anti-CD45-Alexa647-stained CAR T cell interacting with CFSE-labelled HER2+ SKBR3 tumor cell by LLS imaging shows stable cell:cell interaction. Maximum intensity projection from Imaris is shown. Scale bar = 3 μm B. Additional example of a side-view of CAR T cell synapse with HER2+ SKBR3 target prepared as in Fig. 3A-C but with intensity based off endogenous mEmerald tag alone, without additional anti-MYC labelling. CAR localization without anti-MYC labelling shows the same enrichment of CAR in synaptic projections. Maximum intensity projection from Imaris is shown. Scale bar = 2 μm. C. The percentage of microvillar close contacts occupied by receptor was not significantly different between the OT-I TCR and the low-affinity CAR murine T cell synapses with cognate antigen (SL8:pMHC and 625 ng/well HER2). n = 4 cells. D-G. Additional examples of cells prepared and analyzed by line scan as in Fig. 3F-I. H. anti-HER2-BV605 staining was performed on the human breast cancer cell lines MCF7 and SKBR3, plus murine MC38 cells as a HER2-negative control. Histogram overlay shows low and high HER2 expression for MCF7 and SKBR3, respectively. I. Lipid bilayers with varying HER2 densities were built on 5 μm microspheres as standards for comparison to high-HER2 expressing SKBR3 and low-HER2 expressing MCF7s. Representative histogram from two independent experiments. J. Fura-2 ratio images showing calcium flux (yellow) with tracks for interactions of CAR T cells with HER2-bilayers across antigen density and CAR affinity. Control well with no HER2 is shown on left. Images shown are at 3 min following addition of T cells. Scale bars are 7 μm. K. Fura-2 ratio imaging shows calcium flux across affinity and HER2 density, with low-affinity CAR T cells outperforming high-affinity CAR T cells on high-HER2 loaded bilayers. Top: Maximum Fura-2 ratio for cells was quantified at 5.5 min following addition of T cells. Error bars represent s.d. (n = 37, 23, 21, 33, 66 cells per condition from left to right, respectively). Analyses are unpaired t tests. Bottom: Percentage of cells fluxing (defined as maximum Fura-2 ratio > 100) for each condition was quantified at 5.5 min following addition of T cells. L. Schematic of original anti-HER2 CAR, including two cysteines in the CD8 hinge region that form disulfide bond, and monomeric CAR made by mutation of those cysteines to serine hinge (Hennecke and Cosson, 1993). M. The average close contact CAR-mEmerald fluorescence intensity was not significantly different between monomers and dimers. Data is shown for at least 5 cells per condition across three experiments (n = 7, 7, 5, 13 cells per group from left to right, respectively). Error bars represent s.d., analyses shown are unpaired t tests. N. Average close contact CAR-mEmerald fluorescence intensity was plotted against persistence time following color schema in H. Linear trend lines were added for each CAR type (monomers dashed).

**Supplemental Figure 3.**
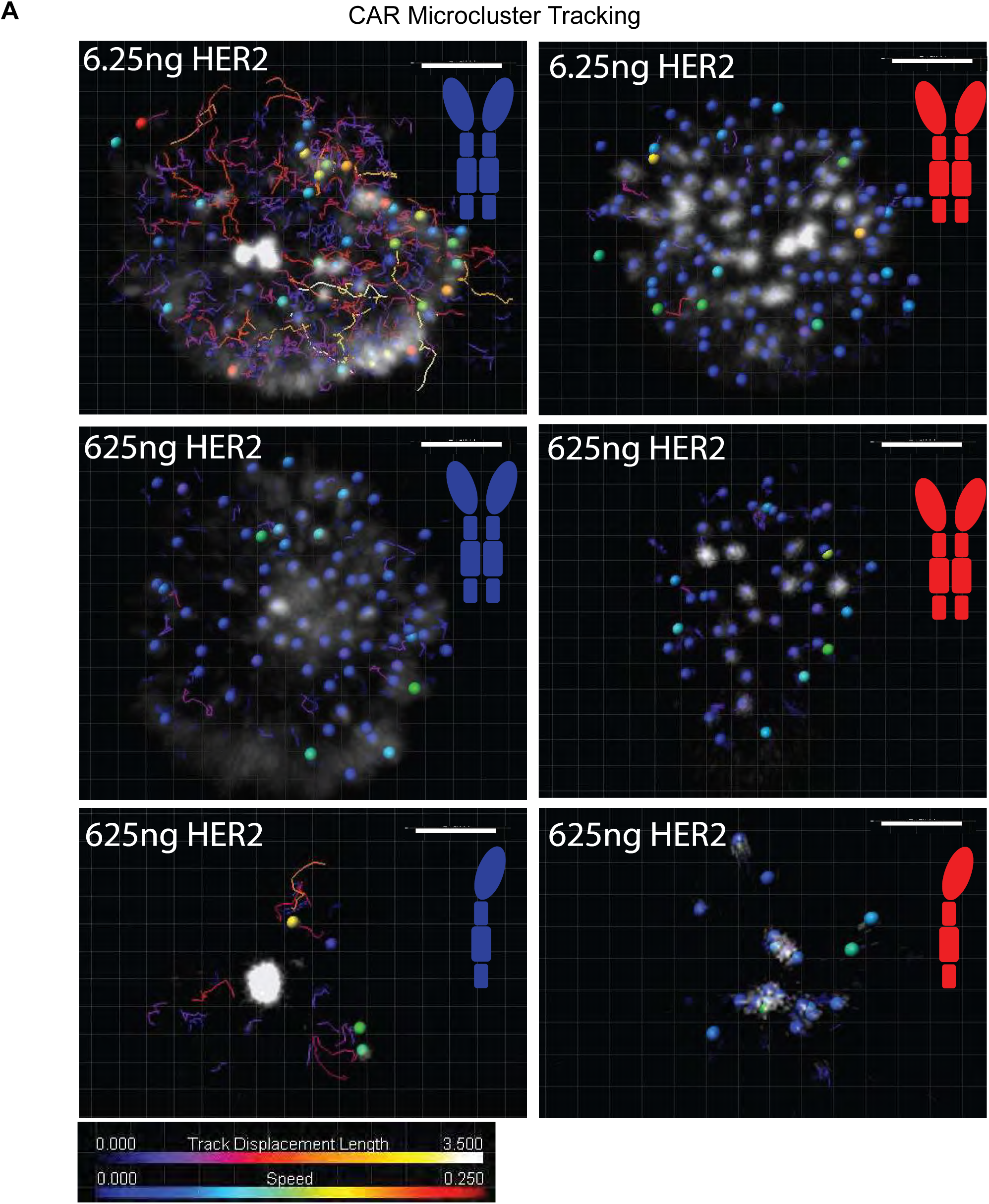
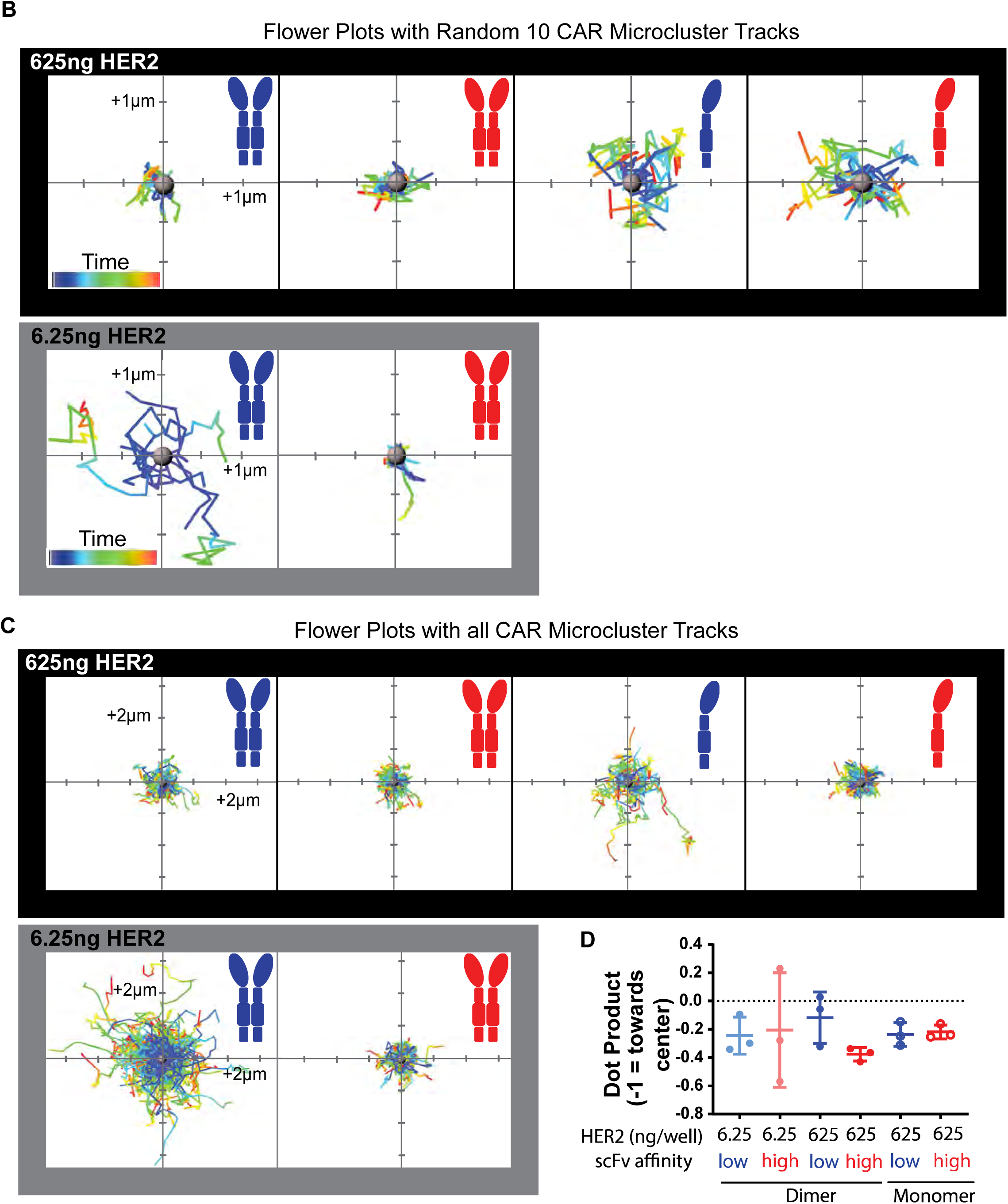
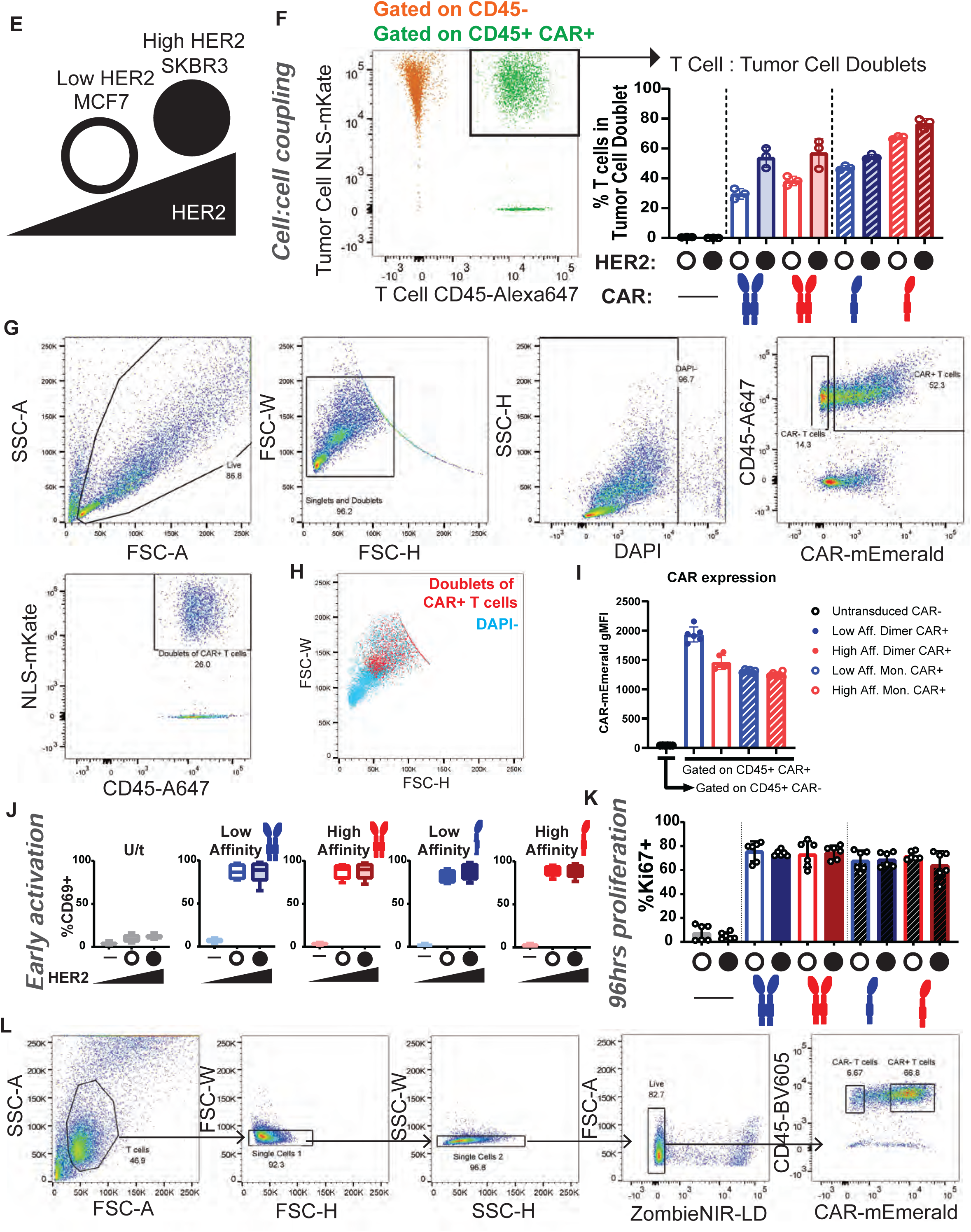
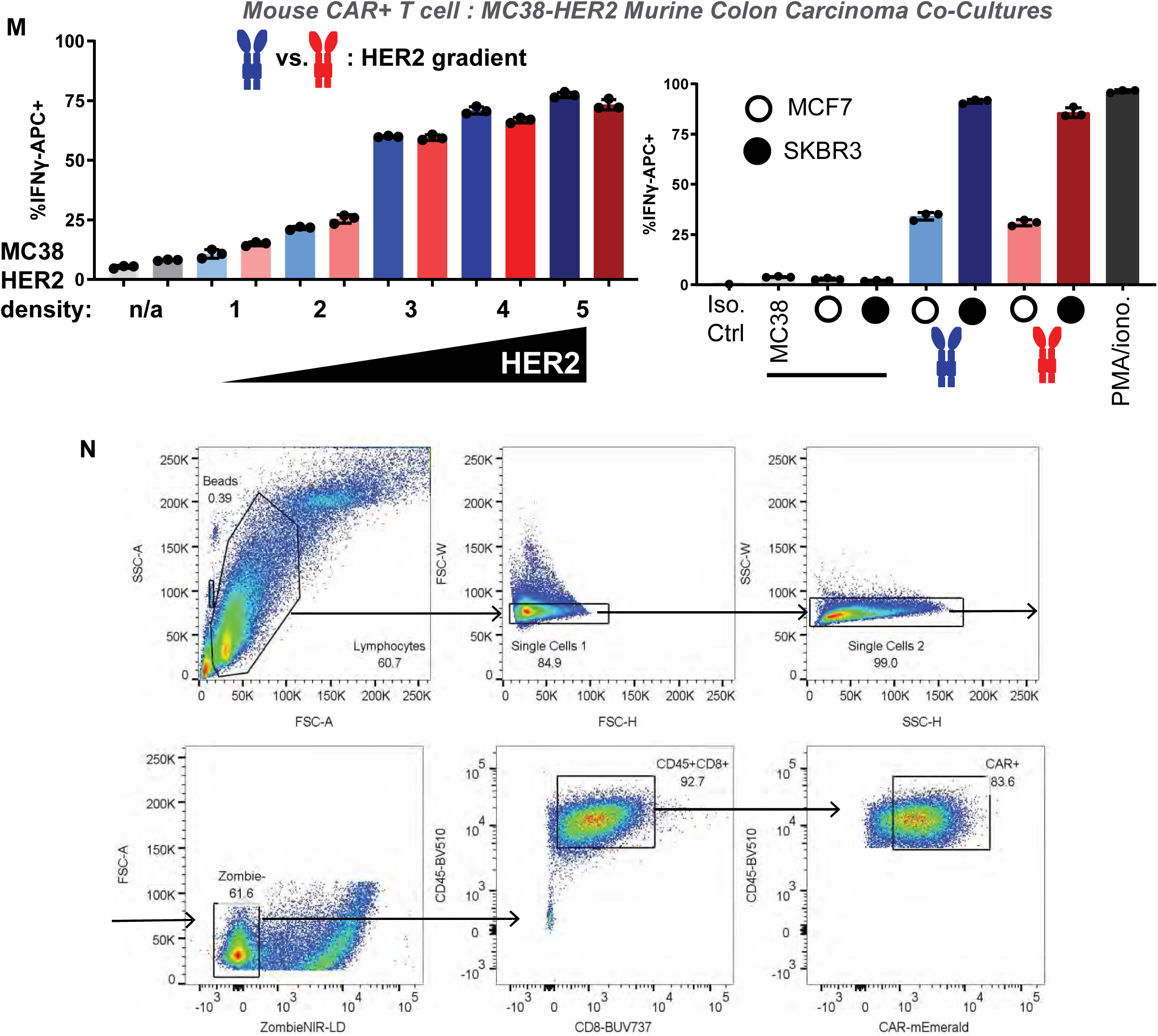
CAR microcluster dynamics and T cell activity. A. TIRF imaging of CAR-mEmerald is shown with spots and tracks made in Imaris. Tracks analysis shows that high-affinity CAR microclusters on low HER2 (6.25 ng/well) are not mobile, and on high HER2 (625 ng/well) only low-affinity monomer is mobile. Scale bars = 3 μm. B. A random 10 tracks are shown in flower plots for each condition. Top: Limited mobility is apparent for dimeric CARs on high-HER2 loaded bilayers. Bottom: On low-HER2 bilayers, low- affinity CAR microclusters show increased mobility. Plots of low affinity dimer from Fig. 5c are shown again here for comparison. C. Flower plots are shown including all tracks for each cell. D. The normalized dot product of V(rad) and V(i) was calculated for all tracks, where V(rad) = vector from synapse center to track start position and V(i) = vector from track start to track final position. Using this calculation, movements directly towards the center equal -1 and movements directly away from center equal 1. The average dot product for all tracks in a cell are quantified. No significant differences are identified in the direction of CAR microcluster movement across conditions. All n.s. by Tukey’s multiple comparisons test. E. Low HER2-expressing MCF7 (open circle) and high HER2-expressing SKBR3 (solid circle) cell lines were used to assess differences between HER2 levels in cell-cell interactions in vitro. F. MCF7 and SKBR3 cells expressing nuclear localization signal (NLS)-tagged mKate were incubated with anti-CD45-Alexa647-labelled T cells for 30 minutes. Coupling analysis was performed by gating inclusively for singlets and doublets. Cells were then gated on DAPI-, CD45+, and CAR+ (or CAR- for untransduced controls). Percentage of these cells that are mKate+ was reported as T cells in doublets with tumor cells (right). Representative dot plot for the high affinity monomeric CAR T cell / SKBR3 co-culture is shown overlaying the CD45- and CD45+CAR+ populations (left). Representative of three independent experiments is shown. G. Gating strategy for coupling assay in (B). Example data here is from a co-culture of low- affinity dimeric CAR T cells with MCF7 target cells. H. The entire DAPI- population in (E) is overlaid with the Doublets of CAR+ T cells population. Plotting by FSC-H vs FSC-W shows that the Doublets of CAR+ T cells population falls in the expected region (above the singlets). I. CAR gMFIs are shown for each receptor type. While there is some variability in expression, differences are within ∼2/3rds of the highest gMFI. J. Early activation is similar across all CAR+ conditions with low and high HER2 at 18 hours following co-incubation, as seen by anti-CD69-BUV395 positivity. Box and whiskers plot error bars showing minimum and maximum values (n = 12 samples per group pooled from four independent experiments and two donors). K. Percentage of anti-Ki67-PE-eFluor 610 positive cells, indicating entry into cell cycle, is similar across all CAR+ conditions with low and high HER2. Replicates from 2 independent experiments of different donors are pooled (n = 6). Error bars represent s.d. L. Gating strategy used for defining human CAR+ (and CAR-) T cells in flow cytometry experiments is shown. Example data shown here is from a co-culture of low-affinity dimeric CAR T cells with SKBR3 target cells. M. Low-affinity (blue) and high-affinity (red) CARs were retrovirally expressed in primary mouse CD8+ T cells. HER2-expressing MC38 cells were sorted into 5 bins of expression levels. MC38- HER2 (1-5) and parental MC38 (n/a) were cultured with CAR T cells for 18 hours and then stained for IFN-γ. Low- and high-affinity CARs performed similarly. Isotype control, untransduced T cell, MCF7 (open circle), SKBR3 (filled circle), and PMA/ionomycin controls are shown on the right. n = 3 replicates per group. Data shown is representative of 3 independent experiments. N. Gating strategy used for defining mouse CAR+ T cells in flow cytometry experiments following in vitro co-culture is shown. Example data shown here is from a co-culture of low- affinity dimeric CAR T cells with MC38-HER2-high targets in normoxia.

## Supplemental Video Captions

**Supplemental Video 1. LLS imaging of anti-HER2 CAR T cell.** Maximum intensity projection of anti-HER2 CAR-expressing T cell shown in Fig. 1a. Location of coverslip, where signal intensity is low, is annotated in yellow. Anti-CD45-Alexa594, anti-MYC (CAR)-Alexa488 and TCR (OKT3)-APC signal is shown in cyan, green and red, respectively.

**Supplemental Video 2. LLS live cell imaging of anti-HER2 CAR T cell in synapse with HER2+ SKBR3 cell.** The blended view of the Imaris volume is shown for anti-HER2 CAR- expressing T cell shown in Fig. 3a. The SKBR3 target cell is located above, as annotated in Fig. 3a. Anti-CD45-Alexa647 and anti-MYC (CAR)-Alexa488 signals are shown in red and green, respectively. Scale bar = 2 μm. Time resolution per frame is 4.7 secs and total video is 20 frames (89.3 secs).

**Supplemental Video 3. Individual close contacts imaged by SCM TIRF microscopy.** QD605 signal is shown for one field of view for low-affinity (top) and high-affinity (bottom) CAR T cells interacting with HER2-loaded bilayer. Holes in QD605 signal show location of microvillar close contacts. High-affinity close contact stably persists in same field of view, while low-affinity CAR T cell contacts appear and disappear from field of view throughout imaging. Scale bar is 0.2 μm. Time resolution per frame is 2.4 secs and total video is 40 frames (93.6 secs).

**Supplemental Video 4. Fura-2 calcium flux imaging of low- and high-affinity CAR T cells on low- and high-HER2 bilayers.** Fura-2 340nm/380nm ratio channel with tracking is shown for A) low affinity/0 ng HER2, B) low affinity/62.5 ng HER2, C) low affinity/625 ng HER2, D) high affinity/62.5 ng HER2, and E) high affinity/625 ng HER2 (from top to bottom) CAR T cells interacting with loaded bilayers. Calcium flux (yellow) is seen for interactions of CAR T cells with HER2-bilayers across antigen density and CAR affinity, relative to no HER2 control (A). Scale bar is 7 μm. Imaging was initiated at 3 min following addition of T cells to wells. Time resolution per frame is 1.5 secs and total video is 100 frames (148.5 secs).

**Supplemental Video 5. CAR microcluster tracking analysis – low-affinity dimer on low HER2 bilayer.** TIRF imaging of CAR-mEmerald is shown for low-affinity dimeric CAR on bilayer loaded with 6.25 ng/well. Spots show position of microclusters used for tracking. Scale bar is 3 μm. Time resolution per frame is 2.4 secs and total video is 40 frames (93.6 secs).

**Supplemental Video 6. CAR microcluster tracking analysis – high-affinity dimer on low HER2 bilayer.** TIRF imaging of CAR-mEmerald is shown for high-affinity dimeric CAR on bilayer loaded with 6.25 ng/well. Spots show position of microclusters used for tracking. Scale bar is 3 μm. Time resolution per frame is 2.4 secs and total video is 40 frames (93.6 secs).

**Supplemental Video 7. CAR microcluster tracking analysis – low-affinity dimer on high HER2 bilayer.** TIRF imaging of CAR-mEmerald is shown for low-affinity dimeric CAR on bilayer loaded with 625 ng/well. Spots show position of microclusters used for tracking. Scale bar is 3 μm. Time resolution per frame is 2.4 secs and total video is 40 frames (93.6 secs).

**Supplemental Video 8. CAR microcluster tracking analysis – high-affinity dimer on high HER2 bilayer.** TIRF imaging of CAR-mEmerald is shown for high-affinity dimeric CAR on bilayer loaded with 625 ng/well. Spots show position of microclusters used for tracking. Scale bar is 3 μm. Time resolution per frame is 2.4 secs and total video is 40 frames (93.6 secs).

**Supplemental Video 9. CAR microcluster tracking analysis – low-affinity monomer on high HER2 bilayer.** TIRF imaging of CAR-mEmerald is shown for low-affinity monomeric CAR on bilayer loaded with 625 ng/well. Spots show position of microclusters used for tracking. Scale bar is 3 μm. Time resolution per frame is 2.4 secs and total video is 40 frames (93.6 secs).

**Supplemental Video 10. CAR microcluster tracking analysis – high-affinity monomer on high HER2 bilayer.** TIRF imaging of CAR-mEmerald is shown for high-affinity monomeric CAR on bilayer loaded with 625 ng/well. Spots show position of microclusters used for tracking. Scale bar is 3 μm. Time resolution per frame is 2.4 secs and total video is 40 frames (93.6 secs).

**Supplemental Table 1.**
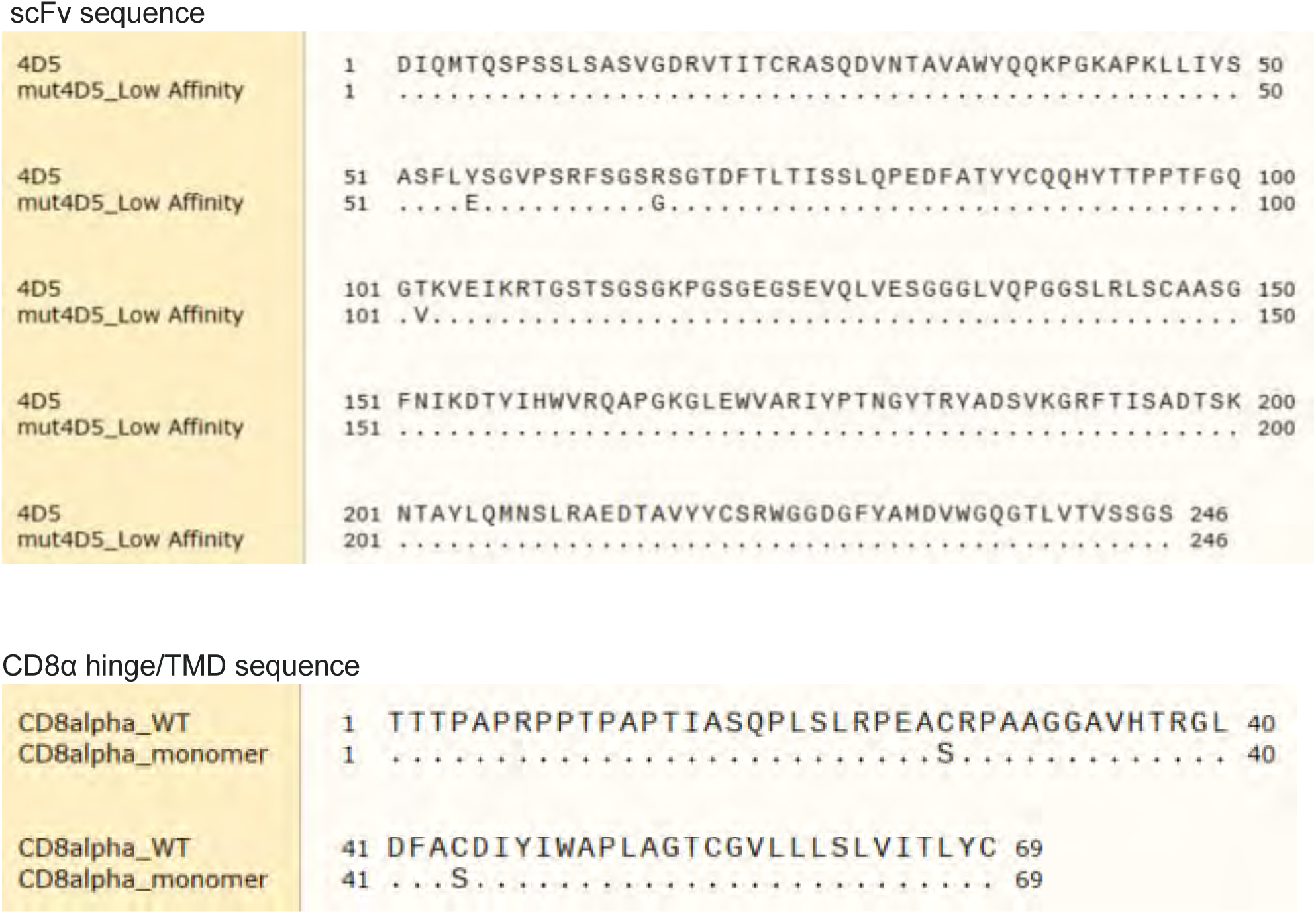
Mutations in CAR scFv (mutCD45) and CD8α hinge/TMD (monomer).

**Supplemental Table 2.** Antibodies referenced for flow cytometry and imaging experiments.

| Anti- | Clone | Fluorophore | Company | Cat. No. |
| --- | --- | --- | --- | --- |
| Myc-Tag | 9B11 | Alexa Fluor 488 | Cell Signaling Tech. | 2233S |
| human CD3 | OKT3 | APC | BioLegend | 317318 |
| human CD340 (HER2) | 24D2 | Brilliant Violet 605 | BD Biosciences | 747615 |
| human CD340 (HER2) | 24D2 | Alexa Fluor 647 | BioLegend | 324412 |
| human CD340 (HER2) | 24D2 | PerCP/Cy5.5 | BioLegend | 324416 |
| human CD45 | HI30 | Alexa Fluor 594 | BioLegend | 304060 |
| human CD45 | HI30 | Alexa Fluor 647 | BioLegend | 304056 |
| human CD45 | HI30 | Brilliant Violet 605 | BioLegend | 304042 |
| human CD69 | FN50 | BUV395 | BD Biosciences | 564364 |
| human IFN-γ | 4S.B3 | APC | BioLegend | 502512 |
| human/mouse Ki67 | SolA15 | PE-eFluor 610 | eBioscience | 61-5698-82 |
| mouse CD223 (LAG3) | C9B7W | Brilliant Violet 785 | BioLegend | 125219 |
| mouse CD279 (PD-1) | 29F.1A12 | Brilliant Violet 421 | BioLegend | 135221 |
| mouse CD279 (PD-1) | 29F.1A12 | Brilliant Violet 605 | BioLegend | 135220 |
| mouse CD39 | Duha59 | PE-Cy7 | BioLegend | 143806 |
| mouse CD4 | GK1.5 | BUV395 | BD Biosciences | 563790 |
| mouse CD44 | IM7 | BUV737 | BD Biosciences | 564392 |
| mouse CD45 | 30-F11 | Brilliant Violet 421 | BioLegend | 103134 |
| mouse CD45 | 30-F11 | Brilliant Violet 510 | BioLegend | 103138 |
| mouse CD45 | 30-F11 | Brilliant Violet 785 | BioLegend | 103149 |
| mouse CD45.1 | A20 | PE-Cy7 | BioLegend | 110730 |
| mouse CD62L | MEL-14 | APC | BioLegend | 104411 |
| mouse CD69 | H1.2F3 | BUV395 | BD Biosciences | 740220 |
| mouse CD69 | H1.2F3 | Brilliant Violet 650 | BioLegend | 104541 |
| mouse CD8a | 53-6.7 | BUV737 | BD Biosciences | 564297 |
| mouse CD8a | 53-6.7 | PerCP/Cy5.5 | BioLegend | 100734 |
| mouse CD90.2 | 30-H12 | Alexa Fluor 700 | BioLegend | 105320 |
| mouse IFN-γ | XMG1.2 | APC | BioLegend | 505810 |
| mouse NK1.1 | PK136 | Brilliant Violet 785 | BioLegend | 108749 |
| mouse TCR $\beta$ chain | H57-597 | Alexa Fluor 488 | BioLegend | 109215 |
| mouse/human CD11b | M1/70 | Brilliant Violet 785 | BioLegend | 101243 |
| mouse/human CD324 (E-cadherin) | DECMA-1 | PE | BioLegend | 147303 |
| mouse/human CD45R | RA3-6B2 | Brilliant Violet 785 | BioLegend | 103246 |
| mouse/human TOX | REA473 | APC | Miltenyi Biotec | 130-118-335 |

## References

Aramesh, M., D. Stoycheva, I. Sandu, S.J. Ihle, T. Zünd, J.Y. Shiu, C. Forró, M. Asghari, M. Bernero, S. Lickert, A. Oxenius, V. Vogel, and E. Klotzsch. 2021. Nanoconfinement of microvilli alters gene expression and boosts T cell activation. Proc. Natl. Acad. Sci. U. S. A. 118. doi:10.1073/pnas.2107535118.

Beemiller, P., J. Jacobelli, and M. Krummel. 2012a. Imaging and Analysis of OT1 T Cell Activation on Lipid Bilayers. Protoc. Exch. doi:10.1038/protex.2012.028.

Beemiller, P., J. Jacobelli, and M.F. Krummel. 2012b. Integration of the movement of signaling microclusters with cellular motility in immunological synapses. Nat. Immunol. 13:787–795. doi:10.1038/ni.2364.

Blumenthal, D., and J.K. Burkhardt. 2020. Multiple actin networks coordinate mechanotransduction at the immunological synapse. J. Cell Biol. 219:1–12. doi:10.1083/jcb.201911058.

Brentjens, R.J., M.L. Davila, I. Riviere, J. Park, X. Wang, L.G. Cowell, S. Bartido, J. Stefanski, C. Taylor, M. Olszewska, O. Borquez-Ojeda, J. Qu, T. Wasielewska, Q. He, Y. Bernal, I. V. Rijo, C. Hedvat, R. Kobos, K. Curran, P. Steinherz, J. Jurcic, T. Rosenblat, P. Maslak, M. Frattini, and M. Sadelain. 2013. CD19-targeted T cells rapidly induce molecular remissions in adults with chemotherapy-refractory acute lymphoblastic leukemia. Sci. Transl. Med. 5. doi:10.1126/scitranslmed.3005930.

Cai, E., C. Beppler, J. Eichorst, K. Marchuk, and M.F. Krummel. 2022. T cells use distinct topological and membrane receptor scanning strategies that individually coalesce during receptor recognition. bioRxiv. 2022.02.23.481517. doi:10.1101/2022.02.23.481517.

Cai, E., K. Marchuk, P. Beemiller, C. Beppler, M.G. Rubashkin, V.M. Weaver, A. Gerard, T.-L. Liu, B.-C. Chen, E. Betzig, F. Bartumeus, and M.F. Krummel. 2017. Visualizing dynamic microvillar search and stabilization during ligand detection by T cells. Science (80-.). 356. doi:10.1126/science.aal3118.

Campi, G., R. Varma, and M.L. Dustin. 2005. Actin and agonist MHC-peptide complex- dependent T cell receptor microclusters as scaffolds for signaling. J. Exp. Med. 202:1031– 1036. doi:10.1084/jem.20051182.

Colin-York, H., Y. Javanmardi, M. Skamrahl, S. Kumari, V.T. Chang, S. Khuon, A. Taylor, T.L. Chew, E. Betzig, E. Moeendarbary, V. Cerundolo, C. Eggeling, and M. Fritzsche. 2019. Cytoskeletal Control of Antigen-Dependent T Cell Activation. Cell Rep. 26:3369–3379.e5. doi:10.1016/j.celrep.2019.02.074.

Comsa, Ş., A.M. Cimpean, and M. Raica. 2015. The Story of MCF-7 Breast Cancer Cell Line: 40 years of Experience in Research. Anticancer Res. 35:3147 LP – 3154.

Corse, E., R.A. Gottschalk, M. Krogsgaard, and J.P. Allison. 2010. Attenuated T cell responses to a high-potency ligand in vivo. PLoS Biol. 8:1–12. doi:10.1371/journal.pbio.1000481.

Crawley, S.W., M.S. Mooseker, and M.J. Tyska. 2014. Shaping the intestinal brush border. J. Cell Biol. 207:441–451. doi:10.1083/jcb.201407015.

Davenport, A.J., R.S. Cross, K.A. Watson, Y. Liao, W. Shi, H.M. Prince, P.A. Beavis, J.A. Trapani, M.H. Kershaw, D.S. Ritchie, P.K. Darcy, P.J. Neeson, and M.R. Jenkins. 2018. Chimeric antigen receptor T cells form nonclassical and potent immune synapses driving rapid cytotoxicity. Proc. Natl. Acad. Sci. U. S. A. 115:E2068–E2076. doi:10.1073/pnas.1716266115.

Dupré, L., K. Boztug, and L. Pfajfer. 2021. Actin Dynamics at the T Cell Synapse as Revealed by Immune-Related Actinopathies. Front. Cell Dev. Biol. 9. doi:10.3389/fcell.2021.665519.

Dushek, O., M. Aleksic, R.J. Wheeler, H. Zhang, S.P. Cordoba, Y.C. Peng, J.L. Chen, V. Cerundolo, T. Dong, D. Coombs, and P.A. Van Der Merwe. 2011. Antigen potency and maximal efficacy reveal a mechanism of efficient T cell activation. Sci. Signal. 4:1–9. doi:10.1126/scisignal.2001430.

Farrell, M. V., S. Webster, K. Gaus, and J. Goyette. 2020. T Cell Membrane Heterogeneity Aids Antigen Recognition and T Cell Activation. Front. Cell Dev. Biol. 8:1–9. doi:10.3389/fcell.2020.00609.

Friedl, P., A.T. Den Boer, and M. Gunzer. 2005. Tuning immune responses: Diversity and adaptation of the immunological synapse. Nat. Rev. Immunol. 5:532–545. doi:10.1038/nri1647.

Friedman, R.S., P. Beemiller, C.M. Sorensen, J. Jacobelli, and M.F. Krummel. 2010. Real-time analysis of T cell receptors in naive cells in vitro and in vivo reveals flexibility in synapse and signaling dynamics. J Exp Med. 207:2733–2749. doi:10.1084/jem.20091201.

Friedman, R.S., J. Jacobelli, and M.F. Krummel. 2006. Surface-bound chemokines capture and prime T cells for synapse formation. Nat. Immunol. 7:1101–1108. doi:10.1038/ni1384.

Garcia, K.C., C.A. Scott, A. Brunmark, F.R. Carbonet, P.A. Peterson, I.A. Wilson, and L. Teyton. 1996. CD8 enhances formation of stable T-cell receptor/MHC class I molecule complexes. 384:577–581.

Ghorashian, S., A.M. Kramer, S. Onuoha, G. Wright, J. Bartram, R. Richardson, S.J. Albon, J. Casanovas-Company, F. Castro, B. Popova, K. Villanueva, J. Yeung, W. Vetharoy, A. Guvenel, P.A. Wawrzyniecka, L. Mekkaoui, G.W.-K. Cheung, D. Pinner, J. Chu, G. Lucchini, J. Silva, O. Ciocarlie, A. Lazareva, S. Inglott, K.C. Gilmour, G. Ahsan, M. Ferrari, S. Manzoor, K. Champion, T. Brooks, A. Lopes, A. Hackshaw, F. Farzaneh, R. Chiesa, K. Rao, D. Bonney, S. Samarasinghe, N. Goulden, A. Vora, P. Veys, R. Hough, R. Wynn, M.A. Pule, and P.J. Amrolia. 2019. Enhanced CAR T cell expansion and prolonged persistence in pediatric patients with ALL treated with a low-affinity CD19 CAR. Nat. Med. 25:1408–1414. doi:10.1038/s41591-019-0549-5.

Grakoui, A., S.K. Bromley, C. Sumen, M.M. Davis, A.S. Shaw, P.M. Allen, and M.L. Dustin. 1999. The Immunological Synapse : A Molecular Machine Controlling T Cell Activation. 221. doi:10.1126/science.285.5425.221.

Gudipati, V., J. Rydzek, I. Doel-Perez, V.D.R. Gonçalves, L. Scharf, S. Königsberger, E. Lobner, R. Kunert, H. Einsele, H. Stockinger, M. Hudecek, and J.B. Huppa. 2020. Inefficient CAR-proximal signaling blunts antigen sensitivity. Nat. Immunol. 21:848–856. doi:10.1038/s41590-020-0719-0.

Harris, D.T., M. V. Hager, S.N. Smith, Q. Cai, J.D. Stone, P. Kruger, M. Lever, O. Dushek, T.M. Schmitt, P.D. Greenberg, and D.M. Kranz. 2018. Comparison of T Cell Activities Mediated by Human TCRs and CARs That Use the Same Recognition Domains. J. Immunol. 200:1088–1100. doi:10.4049/jimmunol.1700236.

Hebeisen, M., L. Baitsch, D. Presotto, P. Baumgaertner, P. Romero, O. Michielin, D.E. Speiser, and N. Rufer. 2013. SHP-1 phosphatase activity counteracts increased T cell receptor affinity. J. Clin. Invest. 123:1044–1065. doi:10.1172/JCI65325.

Hennecke, S., and P. Cosson. 1993. Role of transmembrane domains in assembly and intracellular transport of the CD8 molecule. J. Biol. Chem. 268:26607–26612.

Hu, Y.S., H. Cang, and B.F. Lillemeier. 2016. Superresolution imaging reveals nanometer- and micrometer-scale spatial distributions of T-cell receptors in lymph nodes. Proc. Natl. Acad. Sci. U. S. A. 113:7201–7206. doi:10.1073/pnas.1512331113.

Hudrisier, D., B. Kessler, S. Valitutti, C. Horvath, J.C. Cerottini, and I.F. Luescher. 1998. The efficiency of antigen recognition by CD8+ CTL clones is determined by the frequency of serial TCR engagement. J. Immunol. 161:553–62.

Jung, Y., I. Riven, S.W. Feigelson, E. Kartvelishvily, K. Tohya, M. Miyasaka, R. Alon, and G. Haran. 2016. Three-dimensional localization of T-cell receptors in relation to microvilli using a combination of superresolution microscopies. Proc. Natl. Acad. Sci. U. S. A. 113:E5916– E5924. doi:10.1073/pnas.1605399113.

Kalergis, A.H., N. Boucheron, M.A. Doucey, E. Palmieri, E.C. Goyarts, Z. Vegh, I.F. Luescher, and S.G. Nathenson. 2001. Efficient T cell activation requires an optimal dwell-time of interaction between the TCR and the pMHC complex. Nat. Immunol. 2:229–234. doi:10.1038/85286.

Kalos, M., B.L. Levine, D.L. Porter, S. Katz, S.A. Grupp, A. Bagg, and C.H. June. 2011. T cells with chimeric antigen receptors have potent antitumor effects and can establish memory in patients with advanced leukemia. Sci. Transl. Med. 3:1–13. doi:10.1126/scitranslmed.3002842.

Leithner, A., L.M. Altenburger, R. Hauschild, F.P. Assen, K. Rottner, T.E.B. Stradal, A. Diz- Muñoz, J. V. Stein, and M. Sixt. 2021. Dendritic cell actin dynamics control contact duration and priming efficiency at the immunological synapse. J. Cell Biol. 220. doi:10.1083/JCB.202006081.

Lillemeier, B.F., J.R. Pfeiffer, Z. Surviladze, B.S. Wilson, and M.M. Davis. 2006. Plasma membrane-associated proteins are clustered into islands attached to the cytoskeleton. Proc. Natl. Acad. Sci. U. S. A. 103:18992–18997. doi:10.1073/pnas.0609009103.

Liu, X., S. Jiang, C. Fang, S. Yang, D. Olalere, E.C. Pequignot, A.P. Cogdill, N. Li, M. Ramones, B. Granda, L. Zhou, A. Loew, R.M. Young, C.H. June, and Y. Zhao. 2015. Affinity-tuned ErbB2 or EGFR chimeric antigen receptor T cells exhibit an increased therapeutic index against tumors in mice. Cancer Res. 75:3596–3607. doi:10.1158/0008-5472.CAN-15-0159.

Lyon, A.S., W.B. Peeples, and M.K. Rosen. 2021. A framework for understanding the functions of biomolecular condensates across scales. Nat. Rev. Mol. Cell Biol. 22:215–235. doi:10.1038/s41580-020-00303-z.

Majstoravich, S., J. Zhang, S. Nicholson-Dykstra, S. Linder, W. Friedrich, K.A. Siminovitch, and H.N. Higgs. 2004. Lymphocyte microvilli are dynamic, actin-dependent structures that do not require Wiskott-Aldrich syndrome protein (WASp) for their morphology. Blood. 104:1396–1403. doi:10.1182/blood-2004-02-0437.

Maude, S.L., N. Frey, P.A. Shaw, R. Aplenc, D.M. Barrett, N.J. Bunin, A. Chew, V.E. Gonzalez, Z. Zheng, S.F. Lacey, Y.D. Mahnke, J.J. Melenhorst, S.R. Rheingold, A. Shen, D.T. Teachey, B.L. Levine, C.H. June, D.L. Porter, and S.A. Grupp. 2014. Chimeric Antigen Receptor T Cells for Sustained Remissions in Leukemia. N. Engl. J. Med. 371:1507–1517. doi:10.1056/nejmoa1407222.

McMahan, R.H., J.A. McWilliams, K.R. Jordan, S.W. Dow, D.B. Wilson, and J.E. Slansky. 2006. Relating TCR-peptide-MHC affinity to immunogenicity for the design of tumor vaccines. J. Clin. Invest. 116:2543–2551. doi:10.1172/JCI26936.

Meenderink, L.M., I.M. Gaeta, M.M. Postema, C.S. Cencer, C.R. Chinowsky, E.S. Krystofiak, B.A. Millis, and M.J. Tyska. 2019. Actin Dynamics Drive Microvillar Motility and Clustering during Brush Border Assembly. Dev. Cell. 50:545–556.e4. doi:10.1016/j.devcel.2019.07.008.

Monks, C.R., B. a Freiberg, H. Kupfer, N. Sciaky, and a Kupfer. 1998. Three-dimensional segregation of supramolecular activation clusters in T cells. Nature. 395:82–86. doi:10.1038/25764.

Mossman, K.D., G. Campi, J.T. Groves, and M.L. Dustin. 2005. Altered TCR signaling from geometrically repatterned immunological synapses. Science (80-.). 310:1191–1193. doi:10.1126/science.1119238.

Mota, A. de L., A.F. Evangelista, T. Macedo, R. Oliveira, C. Scapulatempo-Neto, R.A. da C. Vieira, and M.M.C. Marques. 2017. Molecular characterization of breast cancer cell lines by clinical immunohistochemical markers. Oncol. Lett. 13:4708–4712. doi:10.3892/ol.2017.6093.

Nobili, S., A. Mannini, A. Parenti, C. Raggi, A. Lapucci, G. Chiorino, S. Paccosi, P. Di Gennaro, V. Vezzosi, P. Romagnoli, T. Susini, and M. Coronnello. 2021. Establishment and characterization of a new spontaneously immortalized ER−/PR−/HER2+ human breast cancer cell line, DHSF-BR16. Sci. Rep. 11:8340. doi:10.1038/s41598-021-87362-0.

Orbach, R., and X. Su. 2020. Surfing on Membrane Waves: Microvilli, Curved Membranes, and Immune Signaling. Front. Immunol. 11. doi:10.3389/fimmu.2020.02187.

Park, S., E. Shevlin, Y. Vedvyas, M. Zaman, S. Park, Y.M.S. Hsu, I.M. Min, and M.M. Jin. 2017. Micromolar affinity CAR T cells to ICAM-1 achieves rapid tumor elimination while avoiding systemic toxicity. Sci. Rep. 7:1–15. doi:10.1038/s41598-017-14749-3.

Pettmann, J., A.M. Santos, O. Dushek, and S.J. Davis. 2018. Membrane ultrastructure and T cell activation. Front. Immunol. 9:1–9. doi:10.3389/fimmu.2018.02152.

Razvag, Y., Y. Neve-Oz, J. Sajman, M. Reches, and E. Sherman. 2018. Nanoscale kinetic segregation of TCR and CD45 in engaged microvilli facilitates early T cell activation. Nat. Commun. 9. doi:10.1038/s41467-018-03127-w.

Richie, L.I., P.J.R. Ebert, L.C. Wu, M.F. Krummel, J.J.T. Owen, and M.M. Davis. 2002. Imaging synapse formation during thymocyte selection: Inability of CD3ζ to form a stable central accumulation during negative selection. Immunity. 16:595–606. doi:10.1016/S1074-7613(02)00299-6.

Salter, A.I., A. Rajan, J.J. Kennedy, R.G. Ivey, S.A. Shelby, I. Leung, M.L. Templeton, V. Muhunthan, V. Voillet, D. Sommermeyer, J.R. Whiteaker, R. Gottardo, S.L. Veatch, A.G. Paulovich, and S.R. Riddell. 2021. Comparative analysis of TCR and CAR signaling informs CAR designs with superior antigen sensitivity and in vivo function. Sci. Signal. 14. doi:10.1126/scisignal.abe2606.

Sauvanet, C., J. Wayt, T. Pelaseyed, and A. Bretscher. 2015. Structure, Regulation, and Functional Diversity of Microvilli on the Apical Domain of Epithelial Cells. Annu. Rev. Cell Dev. Biol. 31:593–621. doi:10.1146/annurev-cellbio-100814-125234.

Scharping, N.E., D.B. Rivadeneira, A. V. Menk, P.D.A. Vignali, B.R. Ford, N.L. Rittenhouse, R. Peralta, Y. Wang, Y. Wang, K. DePeaux, A.C. Poholek, and G.M. Delgoffe. 2021. Mitochondrial stress induced by continuous stimulation under hypoxia rapidly drives T cell exhaustion. Nat. Immunol. 22:205–215. doi:10.1038/s41590-020-00834-9.

Schmid, D.A., M.B. Irving, V. Posevitz, M. Hebeisen, A. Posevitz-Fejfar, J.-C.F. Sarria, R. Gomez-Eerland, M. Thome, T.N.M. Schumacher, P. Romero, D.E. Speiser, V. Zoete, O. Michielin, and N. Rufer. 2010. Evidence for a TCR Affinity Threshold Delimiting Maximal CD8 T Cell Function. J. Immunol. 184:4936 LP – 4946. doi:10.4049/jimmunol.1000173.

Sendra, G.H., C.H. Hoerth, C. Wunder, and H. Lorenz. 2015. 2D map projections for visualization and quantitative analysis of 3D fluorescence micrographs. Sci. Rep. 5. doi:10.1038/srep12457.

Shakiba, M., P. Zumbo, G. Espinosa-Carrasco, L. Menocal, F. Dündar, S.E. Carson, E.M. Bruno, F.J. Sanchez-Rivera, S.W. Lowe, S. Camara, R.P. Koche, V.P. Reuter, N.D. Socci, B. Whitlock, F. Tamzalit, M. Huse, M.D. Hellmann, D.K. Wells, N.A. Defranoux, D. Betel, M. Philip, and A. Schietinger. 2021. TCR signal strength defines distinct mechanisms of T cell dysfunction and cancer evasion. J. Exp. Med. 219. doi:10.1084/jem.20201966.

Stone, J.D., A.S. Chervin, and D.M. Kranz. 2009. T-cell receptor binding affinities and kinetics: impact on T-cell activity and specificity. Immunology. 126:165–176. doi:10.1111/j.1365-2567.2008.03015.x.

Thompson, S.B., M.M. Waldman, and J. Jacobelli. 2021. Polymerization power: effectors of actin polymerization as regulators of T lymphocyte migration through complex environments. FEBS J. 1–18. doi:10.1111/febs.16130.

Turtle, C.J., L.A. Hanafi, C. Berger, T.A. Gooley, S. Cherian, M. Hudecek, D. Sommermeyer, K. Melville, B. Pender, T.M. Budiarto, E. Robinson, N.N. Steevens, C. Chaney, L. Soma, X. Chen, C. Yeung, B. Wood, D. Li, J. Cao, S. Heimfeld, M.C. Jensen, S.R. Riddell, and D.G. Maloney. 2016. CD19 CAR-T cells of defined CD4+:CD8+ composition in adult B cell ALL patients. J. Clin. Invest. 126:2123–2138. doi:10.1172/JCI85309.

Varma, R., G. Campi, T. Yokosuka, T. Saito, and M.L. Dustin. 2006. T Cell Receptor-Proximal Signals Are Sustained in Peripheral Microclusters and Terminated in the Central Supramolecular Activation Cluster. Immunity. 25:117–127. doi:10.1016/j.immuni.2006.04.010.

Viola, A., S. Linkert, and A. Lanzavecchia. 1997. A T cell receptor (TCR) antagonist competitively inhibits serial TCR triggering by low-affinity ligands, but does not affect triggering by high-affinity anti-CD3 antibodies. Eur. J. Immunol. 27:3080–3083. doi:10.1002/eji.1830271146.

Yokosuka, T., K. Sakata-Sogawa, W. Kobayashi, M. Hiroshima, A. Hashimoto-Tane, M. Tokunaga, M.L. Dustin, and T. Saito. 2005. Newly generated T cell receptor microclusters initiate and sustain T cell activation by recruitment of Zap70 and SLP-76. Nat. Immunol. 6:1253–1262. doi:10.1038/ni1272.

